# Glycosaminoglycans Promote Amyloid-β Aggregation via Multivalent, pH-Dependent Interactions

**DOI:** 10.64898/2026.05.20.726382

**Authors:** Eunjin Moon, Charlotte Radelof, Jana Sticht, Yu Wang, Felicia Fürstenberg, Clemens Krage, Sebastian Straeten, Wesley Pietsch, Boris Schade, Artem Pavlov, Ricardo Zarate, Gaël M. Vos, Gergo Peter Szekeres, Birgit Strodel, Beate Koksch, Kevin Pagel, Nicklas Österlund

## Abstract

Glycosaminoglycans (GAGs) are polyanionic polysaccharides that co-localize with amyloid-β (Aβ) deposits in Alzheimer’s disease, yet their mechanistic contribution to Aβ aggregation remains unclear. Here, we show that GAGs function as pH-responsive electrostatic scaffolds that selectively accelerate Aβ(1–42) aggregation under mildly acidic, endosomal conditions but not at neutral extracellular pH. Combining experimental and computational approaches, we identify protonated N-terminal histidines as key determinants of GAG binding. Weak interactions between GAGs and the charged Nterminal region of Aβ promote conformational rearrangements that bring peptides into proximity and expose adjacent hydrophobic aggregation-prone segments, thereby facilitating peptide clustering. Kinetic analyses reveal that aggregation is enhanced in a way consistent with an apparent increase in effective peptide concentration, accelerating nucleation without altering the dominant aggregation pathway. Systematic variation of GAG chain length and sulfation level further demonstrates that aggregation enhancement requires a threshold degree of multivalency, consistent with a clustering-driven mechanism. Together, these findings establish a framework in which pH-dependent electrostatic interactions with GAGs act as molecular triggers of amyloid nucleation, providing insight into how cellular microenvironments regulate the earliest stages of Alzheimer’s disease pathology.

## Introduction

Glycosaminoglycans (GAGs) are linear anionic polysaccharides that are typically covalently attached to core proteins to form proteoglycans on cell surfaces where they mediate diverse functions, including cell adhesion, signaling, and protein stability and trafficking.^1-3^ GAGs are composed of repeating disaccharide units consisting of a hexuronic acid and a hexosamine and are classified into several families, including heparan sulfate/heparin (HS/Hep), chondroitin sulfate and dermatan sulfate (CS/DS), and hyaluronic acid (HA) (**Figure 1A**).^4^ These classes differ in both composition and degree of sulfation, resulting in distinct charge densities and interaction properties: HS and Hep are highly sulfated, CS and DS are more moderately sulfated, and HA is unsulfated.^5^ The high charge density and structural flexibility of sulfated GAGs enable multivalent interactions with a wide range of proteins, often modulating their conformation and activity.^6-8^ Dysregulation of GAG biosynthesis or turnover has been implicated in numerous pathological conditions, including cancer, fibrosis, inflammatory disorders, and neurodegenerative diseases.^6^ Notably, GAGs are consistently found as integral components of pathological amyloid deposits in aggregation-related diseases, where they co-localize with and can directly interact with aggregation-prone proteins.^9-13^ These pathological interactions may involve either proteoglycans or free GAG chains, such as heparin released during mast cell activation,^14,15^ as well as free HS fragments generated locally by endolysosomal heparanase activity.^16,17^

**Figure 1.**
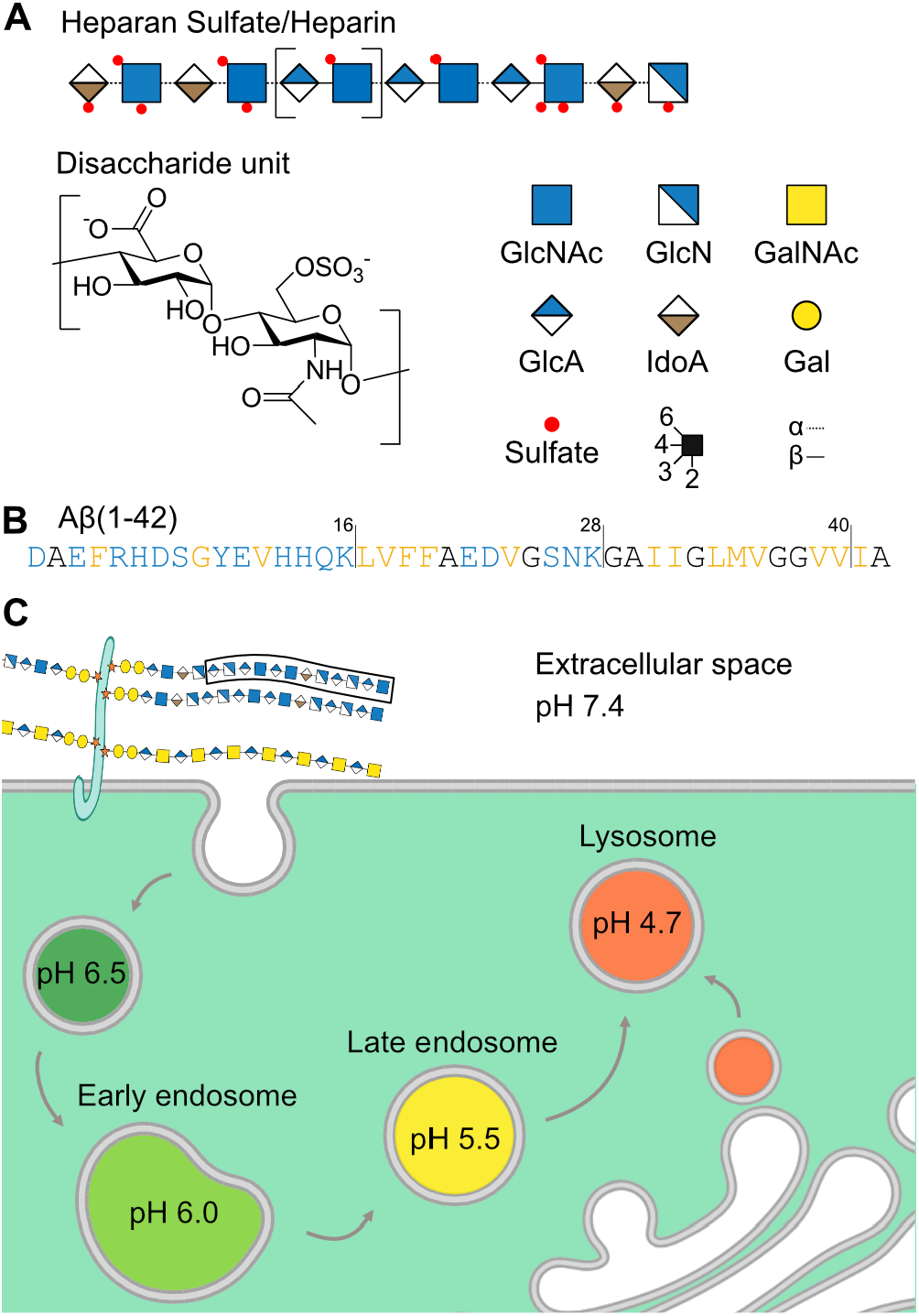
GAG and Aβ peptide properties and the pH microenvironments in cellular compartments. **A)** The GAG disaccharide unit in heparan sulfate is shown as an example along with the SNFG nomenclature for different monosaccharides: *N*-acetylglucosamine (GlcNAc), glucosamine (GlcN), *N*-acetylgalactosamine (GalNAc), glucuronic acid (GlcA), iduronic acid (IdoA), and galactose (Gal). **B)** The sequence of Aβ(1–42) with hydrophobic residues colored orange and hydrophilic residues colored blue. **C)** The cartoon depiction of the cellular microenvironments and their respective pH values.

GAGs have been recognized as important modulators of amyloidogenesis across diverse proteins, affecting aggregation rates or inducing specific fibril polymorphs.^13,18,19^ In Alzheimer’s disease (AD), the amyloid-β peptide is known to interact with GAGs, with which they co-localize in senile plaques.^20,21^ Aβ peptides are generated through proteolytic processing of the membrane-bound amyloid precursor protein (APP) at the plasma membrane or within endosomes,^22,23^ with endosomal cleavage favoring increased Aβ production.^24,25^ Among the different Aβ isoforms, the 42-residue-long Aβ(1–42) variant is particularly prone to self-assembly into β-sheet-rich aggregates associated with neurotoxicity.^26^ Structurally, the peptide consists of a hydrophilic N-terminal region, followed by two hydrophobic segments separated by a flexible hinge region (**Figure 1B**). The aggregation behavior of Aβ is highly sensitive to its microenvironment, including pH,^27^ ionic strength,^28^ and interactions with biological macromolecules.^29-31^

Importantly, Aβ and GAGs co-localize across distinct cellular compartments that differ in physico-chemical conditions. While the extracellular environment is maintained at near-neutral pH, endosomal and lysosomal compartments are mildly acidic (**Figure 1C**) and have been proposed as key sites for Aβ aggregation.^32-38^ GAGs and proteoglycans also facilitate the uptake and trafficking of extracellular Aβ, further linking these environments.^39-41^ Protein–GAG interactions are often mediated by clusters of basic residues,^42,43^ and the three histidine residues in the N-terminal segment of Aβ are particularly relevant in this context due to their pH-sensitive protonation behavior. As the pH decreases from neutral extracellular conditions to mildly acidic endosomal environments, protonation of these residues increases the local positive charge of the peptide, which may strengthen electrostatic interactions with negatively charged GAGs.

While these observations highlight the importance of environmental and molecular factors in peptide– GAG interactions, the underlying mechanisms remain incompletely understood. Previous studies have reported both enhancement and inhibition of Aβ fibril formation in the presence of GAGs.^9,44,45^ These findings may arise from inadequate control over multiple key variables, including differences in GAG class, chain length, sulfation pattern, peptide aggregation state, and solution conditions. For example, different GAG classes can exert distinct biological effects, and aberrant heparan sulfate expression has been observed in patients with AD.^46-48^ In addition, Aβ may interact differently with GAGs depending on its aggregation state, with stronger binding reported for specific aggregated species.^49,50^ These variables complicate the interpretation of GAG-mediated effects and obscure the mechanistic basis of their interaction with Aβ.

Here, we examine well-defined GAG samples to assess their binding to soluble forms of the Aβ peptide and the effects on aggregation of these peptide species. We demonstrate a pronounced pHswitchable behavior of the Aβ peptide that governs its interaction with GAGs via two well-defined binding patches in its N-terminal region. These interactions, in turn, accelerate peptide aggregation by providing favorable sites for peptide clustering. We show that this effect is governed by relatively weak electrostatic interactions, scales with both the GAG chain length and the degree of sulfation, and requires a threshold level of multivalency for the aggregation-enhancing effect.

## Results

### GAGs promote Aβ aggregation at endosomal pH

We first examined the interaction between Aβ and GAGs at pH values representative of the two environments in which these interactions are expected to occur: the extracellular (pH 7.4) and endosomal (pH 6.0) environments. To assess the effect of GAGs, we added enoxaparin, a clinically used low-molecular-weight heparin (LMWH), and monitored the effect on Aβ(1–42) amyloid aggregation. Aggregation was followed by measuring the time-dependent fluorescence of the amyloid-selective probe Thioflavin T (ThT), whose quantum yield increases markedly upon binding to amyloid fibrils.^51^ At pH 7.4, addition of an equimolar amount of LMWH did not alter the ThT fluorescence kinetics compared to the control experiment of Aβ(1–42) alone (**Figure 2A**). In contrast, when the pH was lowered to 6.0, the same amount of LMWH accelerated Aβ aggregation (**Figure 2B**). Both the lag time (*t*_*lag*_) and maximum growth rate (*r*_*max*_) of aggregation were affected, with *t*_*lag*_ decreasing from 43 ± 2 min to 25 ± 0.6 min, and *r*_*max*_ increasing from 2.45 ± 0.25 h^-1^ to 4.8 ± 0.73 h^-1^.

**Figure 2.**
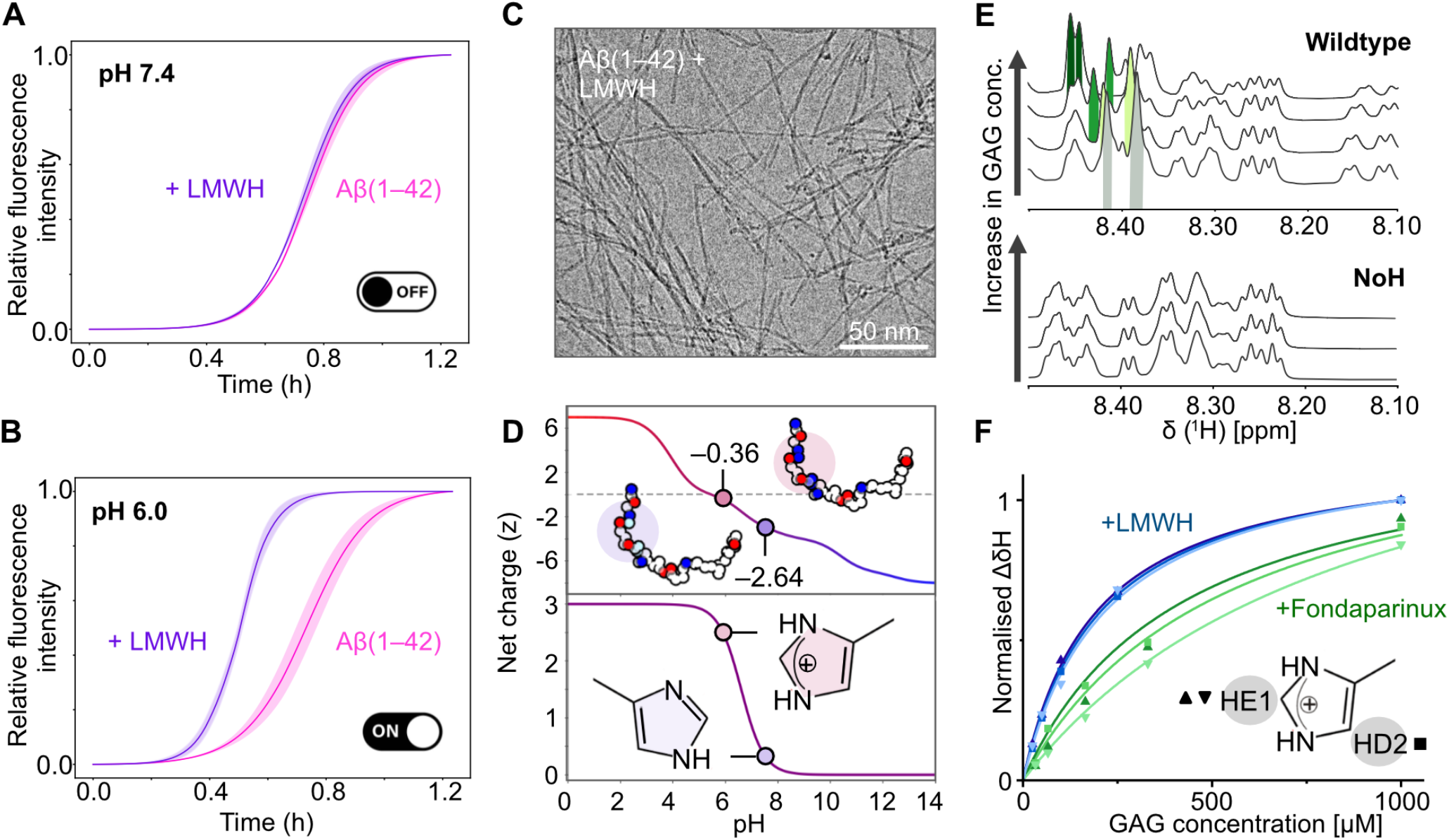
The effect of low-molecular-weight heparin on amyloid aggregation kinetics. Amyloid aggregation of Aβ(1–42) followed by monitoring the fluorescence of ThT in the absence (pink) and presence (purple) of low-molecular weight heparin (LMWH) at **A)** pH 7.4 and **B)** pH 6.0. **C)** Cryo-EM micrograph of Aβ(1–42) fibrils (20 µM) upon incubation with equimolar amounts of LMWH for 2 days at pH 6.0. **D)** Theoretical net charge of Aβ(1–42) (top) and the three histidine residues (bottom) as a function of pH. **E)** ^1^H NMR spectra of Aβ(1–28) and the Aβ(1– 28) H6A/H13A/H14A triple mutant (NoH) upon increasing concentrations of the pentasaccharide fondaparinux at pH 6.0. The resonances corresponding to histidine HE1 protons are highlighted. **F)** Normalized shifts in ^1^H chemical shift difference (ΔδH) in Aβ(1–28) resonances upon titration with LMWH (blue) or fondaparinux (green) at pH 6.0 for HE1 (triangles) and HD2 (squares) resonances.

Control ThT experiments performed with LMWH in the absence of Aβ confirmed that the GAG component itself does not increase ThT fluorescence (**Figure S1**). To verify that the enhanced ThT signal reflected genuine amyloid formation rather than an assay artifact, we complemented the fluorescence measurements with label-free structural and morphological analyses. Cryogenic electron microscopy (cryo-EM) showed that the addition of equimolar amounts of LMWH at pH 6.0 led to the formation of typical long and twisted amyloid fibrils (**Figure 2C**). Some changes in fibril morphology could, however, be observed, with the fibrils formed with LMWH being longer and more curved compared to the control sample without LMWH (**Figure S3**). Circular Dichroism (CD) spectroscopy further confirmed the appearance of a β-sheet-rich structure upon incubation with LMWH at pH 6.0 (**Figure S2**). These findings confirm that the accelerated aggregation observed in the ThT assay still results in amyloid fibril formation, rather than generating more amorphous aggregates, as has been observed in other aggregating systems upon modulating ionic strength.^52^

The observed pH dependence may point to a role for histidine residues in mediating GAG-induced aggregation. Within the investigated pH range, histidines are the primary residues expected to undergo substantial changes in protonation. Lowering the pH from 7.4 to 6.0 increases the combined charge of the three histidine residues from +0.36 to +2.32, shifting the net peptide charge from −2.64 to −0.36 based on experimentally determined pK_a_ values (**Figure 2D**).^53^ This effect is even more pronounced for the N-terminal segment (residues 1–16), which becomes net positively charged at pH 6.0. One possible interpretation is that the increase in local positive charge enhances electrostatic interactions with negatively charged GAG chains, thereby promoting aggregation under mildly acidic conditions.

### Aβ–GAG interactions depend on histidine residues

To test our hypothesis regarding the GAG interaction site in the peptide, we next synthesized model peptides comprising residues 1–28 of Aβ. This segment contains the hydrophilic, pH-sensitive Nterminal region, while removal of the highly hydrophobic C-terminal portion yields a peptide with substantially reduced propensity to aggregate. In this way, we sought to decouple GAG binding from aggregation, enabling an unbiased assessment of the binding mechanism while minimizing interference with peptide aggregation. We synthesized both the wild-type Aβ(1–28) sequence and a histidine-substituted variant in which residues H6, H13, and H14 were replaced by alanine (NoH) (**Figure S4**). Both peptides were C-terminally amidated to avoid introducing a negative charge at position 28, which is not present in the full-length peptide.

To probe peptide–GAG interactions, we employed solution-state NMR spectroscopy. ^1^H-NMR allows direct observation of binding without labeling and can provide residue-specific information about the interaction. To further reduce system complexity, we first used the GAG pentasaccharide fondaparinux, a synthetic heparin mimetic with a well-defined sequence. We recorded 1D ^1^H spectra of Aβ(1– 28) and fondaparinux separately, confirming that their characteristic resonances do not overlap (**Figure S5**). The HE1 and HD2 resonances of the histidine side chain were identified in the 2D TOCSY and ^1^H-^13^C HMQC spectra (**Figure S6**).

We next stepwise titrated the GAGs into the peptide at pH 6.0. Increasing GAG concentrations induced selective shifts in selected proton resonances (**Figure 2E**, top), indicating preferential interaction with specific regions of the peptide. The gradual, concentration-dependent chemical-shift changes are consistent with fast exchange on the NMR timescale, with peak positions reflecting population-weighted averages of the resonance positions of the bound and unbound states, respectively.^54^ In contrast, titration of fondaparinux to the NoH control peptide resulted in no detectable spectral changes (**Figure 2E**, bottom), confirming that histidine residues are essential for GAG binding. Titrations to Aβ(1–28) were next performed using LMWH. The chemical-shift changes of the affected resonances were plotted as a function of GAG concentration for both fondaparinux and LMWH (**Figure 2F**). From these binding curves, we estimated apparent dissociation constants (*K*_*d*_) of 170 ± 40 µM for LMWH and 940 ± 270 µM for fondaparinux, with the latter showing markedly weaker binding. GAG binding at acidic pH was independently confirmed using surface plasmon resonance (SPR) using immobilized LMWH (**Figure S7**).

Upon addition of LMWH, we observed a gradual loss of NMR signal intensity, suggesting that the low-aggregation-propensity Aβ(1–28) peptide may aggregate upon GAG interaction (**Figure S8**). This effect was much weaker in titrations with fondaparinux. To assess such GAG-induced aggregation, Aβ(1–28) and NoH were monitored by CD spectroscopy upon addition of LMWH at pH 6.0. Both peptides initially adopted stable random-coil structures for ∼2 h, consistent with their low intrinsic aggregation propensity. Addition of LMWH triggered structural changes in Aβ(1–28), followed by gradual conversion to a mature β-sheet (**Figure S9A**). In contrast, NoH showed minimal response to LMWH and remained largely disordered throughout incubation (**Figure S9B**).

### GAG binding occurs to two N-terminal binding motifs in Aβ

Next, to obtain per-residue information about the GAG-binding site within the peptide, we conducted HSQC experiments. This experiment provides one cross-peak for each backbone amide, generating a 2D fingerprint spectrum in which individual amino acids are better resolved compared to a onedimensional ^1^H spectrum and can be monitored separately. All measurements were performed on unlabeled peptides, relying on the natural abundance of ^15^N. Aβ(1–28) yielded a well-resolved HSQC spectrum (**Figure 3A**), with cross peaks assigned based on 2D-NOESY, 2D-TOCSY, ^1^H-^13^C-HMQC, and ^1^H-^15^N-HSQC spectra (**Figure S6**), also taking into account previously published assignments and predicted chemical shifts (*see Methods*).^55^ Upon titration of fondaparinux, several peptide resonances exhibited clear chemical-shift perturbations. Quantification of these changes using the combined chemical-shift perturbation metric (Δδ(^1^H, ^15^N)) revealed two distinct regions affected by GAG addition: residues 4–6 and residues 12–17 (**Figure 3B**). These two clusters include the majority of positively charged residues in the peptide, including _12_VHHQKL_17_, which at low pH conforms to the previously suggested heparin-binding motif XBBXBX, where B is a positively charged residue, and X is any other residue. Notably, the basic residue K28, located in the C-terminal region, did not exhibit detectable perturbations upon GAG addition.

**Figure 3.**
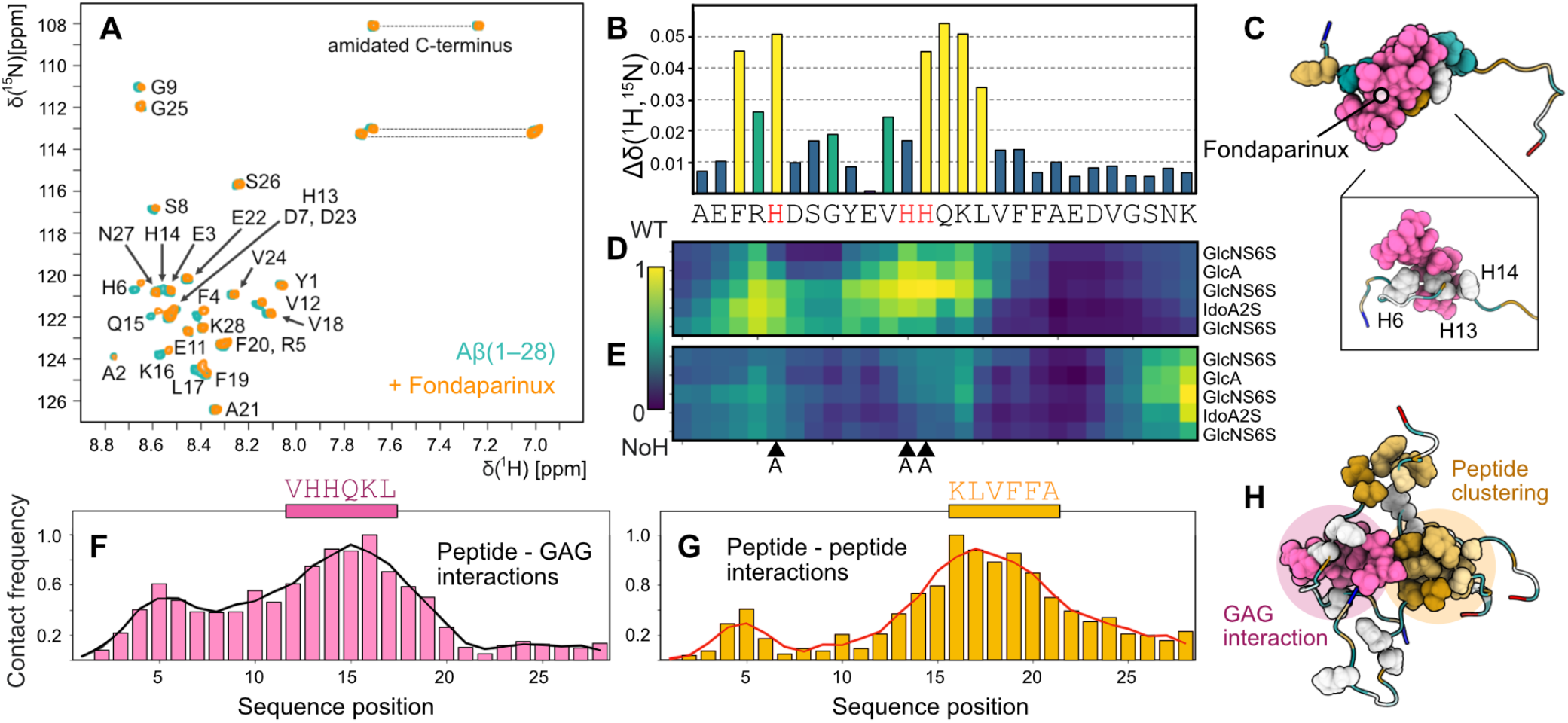
Probing the GAG/Aβ binding site. **A)** ^1^H–^15^N HSQC NMR spectra of Aβ(1–28) before (green) and after (orange) addition of the pentasaccharide fondaparinux. **B)** Combined chemical-shift perturbation from panel A plotted over the sequence of Aβ(1–28), note that residue D1 is not detected. The major shifts (> mean shift + standard deviation) are colored yellow and minor shifts (> mean shift + half of the standard deviation) in green. **C)** Visualization of the top cluster structure of Aβ(1–28) (colored according to hydrophobicity with brown representing hydrophobic residues and blue representing hydrophilic residues) and fondaparinux (pink). The inset shows the interaction between the GAG (pink) and the three histidines of Aβ (gray, spherical representation). **D–E)** Contact map of the peptide–GAG interactions from MD simulations of the Aβ(1–28) (D) and NoH (E) monomers in the presence of one fondaparinux. **F–G)** Relative intermolecular contact frequency between Aβ and fondaparinux (F) and across Aβ peptides (G) in the system comprising three peptides and one GAG molecule. The line represents the running average. **H)** Visualization of the top cluster structure of three Aβ(1–28) (colored according to hydrophobicity) and one fondaparinux molecule (pink). The histidines of Aβ and the central hydrophobic cluster KLVFFA are shown in spherical representation to highlight the distinct peptide–GAG and peptide–peptide interaction hotspots.

To further elucidate the structural basis of the peptide–GAG interaction, we next turned to molecular dynamics (MD) simulations. All-atom MD simulations were performed in triplicate for Aβ(1–28) and NoH monomers with and without fondaparinux at a 1:1 molar ratio. The MD simulations reveal a common binding mode in which the GAG orients approximately perpendicular to the peptide’s N-terminal region (**Figure 3C**). In this arrangement, the central core of the pentasaccharide forms a binding pocket that accommodates the histidine residues of Aβ. Analysis of the conformational ensemble indicates that both the wild-type and NoH variants predominantly populate disordered states. However, the wild-type peptide shows a clear propensity for transient β-hairpin formation, whereas the mutant instead favors α-helical conformations, consistent with the helix-stabilizing effect of the alanine substitution. Upon GAG binding, the wild-type peptide adopts a more extended conformation, reflected by an increased N–C terminal distance, a rearrangement associated with facilitating subsequent interpeptide β-sheet assembly (**Figure S10**). In contrast, the triple mutant peptide exhibits only minor structural perturbations following GAG addition, with infrequent peptide–GAG contacts and a largely unchanged conformational ensemble (**Figure S11**).

Consistent with the binding mode identified above, peptide–GAG contact maps of wild-type Aβ(1–28) in complex with fondaparinux reveal extensive interactions, indicating near-continuous association between the peptide and the ligand (**Figure 3D, Figure S12A**). In excellent agreement with the HSQC experiments, two regions showed strong engagement: residues 4–6 around R5, which displayed similar contact probabilities with nearly all monosaccharide rings of the pentasaccharide, and the central segment comprising residues 10–17, which preferentially interacts with the central sugar ring of the GAG pentasaccharide. These interactions are completely absent in the contact map of the NoH variant with fondaparinux (**Figure 3E**). The total contact frequency of NoH with fondaparinux is only 5% of the contact frequency of the wild-type Aβ(1–28), indicating that the remaining positively charged residues are not enough to rescue the interaction (**Figure S12B**). The highest interactions were confined to K28 and its immediate neighbors (15% contact frequency relative to the wild type), whereas R5 and K16 exhibited very low contact probabilities.

To assess the effect of GAG binding on a system of multiple assembling Aβ peptides, we next performed MD simulations of three wild-type Aβ(1–28) peptides in the presence of a single fondaparinux molecule. The simulations show that the segment _12_VHHQKL_17_ remains the principal interaction site for fondaparinux in this system as well (**Figure 3F**). Notably, one peptide acts as the dominant binding partner for fondaparinux, whereas the remaining peptides engage in more diffuse and transient interactions (**Figure S13**). This behavior is consistent with the preferred GAG interaction site, the central sugar unit of the pentasaccharide, being occupied and thereby limiting equivalent binding by the other peptides. Analysis of intermolecular peptide–peptide contacts reveals a clear preferred interaction hotspot formed by the central hydrophobic cluster _16_KLVFFA_21_, a region well known for its critical role in Aβ aggregation, which engages in self-association between peptides (**Figure 3G, Figure S14**). The same aggregation hotspot is also seen in control simulations of Aβ only, indicating that GAG binding does not substantially alter peptide–peptide clustering **(Figure S14)**.

To experimentally probe the structural interplay between GAG binding and aggregation, we next analyzed the truncated variants Aβ(1–16) and Aβ(12–28) in the presence of LMWH using NMR. The N-terminal Aβ(1–16) fragment, which lacks the hydrophobic, aggregation-prone region, exhibited histidine-dependent fast-exchange binding behavior similar to that of Aβ(1–28), but with reduced line broadening, consistent with diminished aggregation (**Figure S15**). In contrast, Aβ(12–28), which retains the second histidine-rich GAG binding region and the hydrophobic segment, displayed extensive line broadening across the spectrum, indicative of intermediate exchange and/or aggregation (**Figure S16**). Notably, residues distal to the putative GAG binding site were also affected, suggesting that the observed spectral changes in the spectrum for the 12–28 fragment are dominated by aggregation rather than simple binding.

These observations support that histidine-mediated GAG binding and hydrophobic clustering of peptides are intrinsically coupled processes. The N-terminal region binds to GAGs electrostatically, while the adjacent hydrophobic segment promotes peptide–peptide interactions that compete with or mask GAG binding, leading to apparent changes in exchange behavior and affinity as observed by NMR. The peptide–peptide interaction hotspot is positioned immediately after the peptide–GAG binding region (**Figure 3H**), suggesting that GAG association promotes spatial clustering of peptides around the GAG scaffold while simultaneously exposing aggregation-prone segments in its vicinity. This arrangement, which affects both local peptide concentration and peptide conformations, is likely to increase the probability of productive nucleation events during assembly.

### GAGs increase the apparent Aβ concentration during aggregation

To test the idea that the polyanionic GAG chain changes the concentration dependence of aggregation by promoting peptide clustering, we next conducted ThT kinetic experiments at varying Aβ monomer concentrations (*m*_*0*_) with and without LMWH (**Figure 4A–B**). It is well established that the aggregation half-time (*t*_*1/2*_), which scales proportionally with *t*_*lag*_, exhibits a power-law dependence on *m*_*0*_ according to *t*_*1/2*_ =A m_0_^γ^, where γ is the apparent reaction order of the aggregation process, and A is a kinetic prefactor.^56-58^ For aggregation dominated by secondary nucleation, γ is given by γ = −(*n*_*c*_+1)/2, where *n*_*c*_ is the critical nucleus size for secondary nucleation. For Aβ(1–42) at pH 8, γ values between −1.5 and −1 have been reported.^59,60^ In our data at pH 6, we observe γ = −0.47 ± 0.04 for Aβ(1–42) alone (**Figure 4C**). This value, close to γ = −1/2, indicates saturation of the secondary nucleation step (*n*_*c*_ = 0). In this regime, nucleation occurs when the fibril surface is fully occupied. At lower pH, closer to the isoelectric point of the peptide (pI = 5.4), saturation is likely promoted by reduced electrostatic repulsion and more favorable adsorption to the fibril surface compared to pH 8.^27,59^

**Figure 4.**
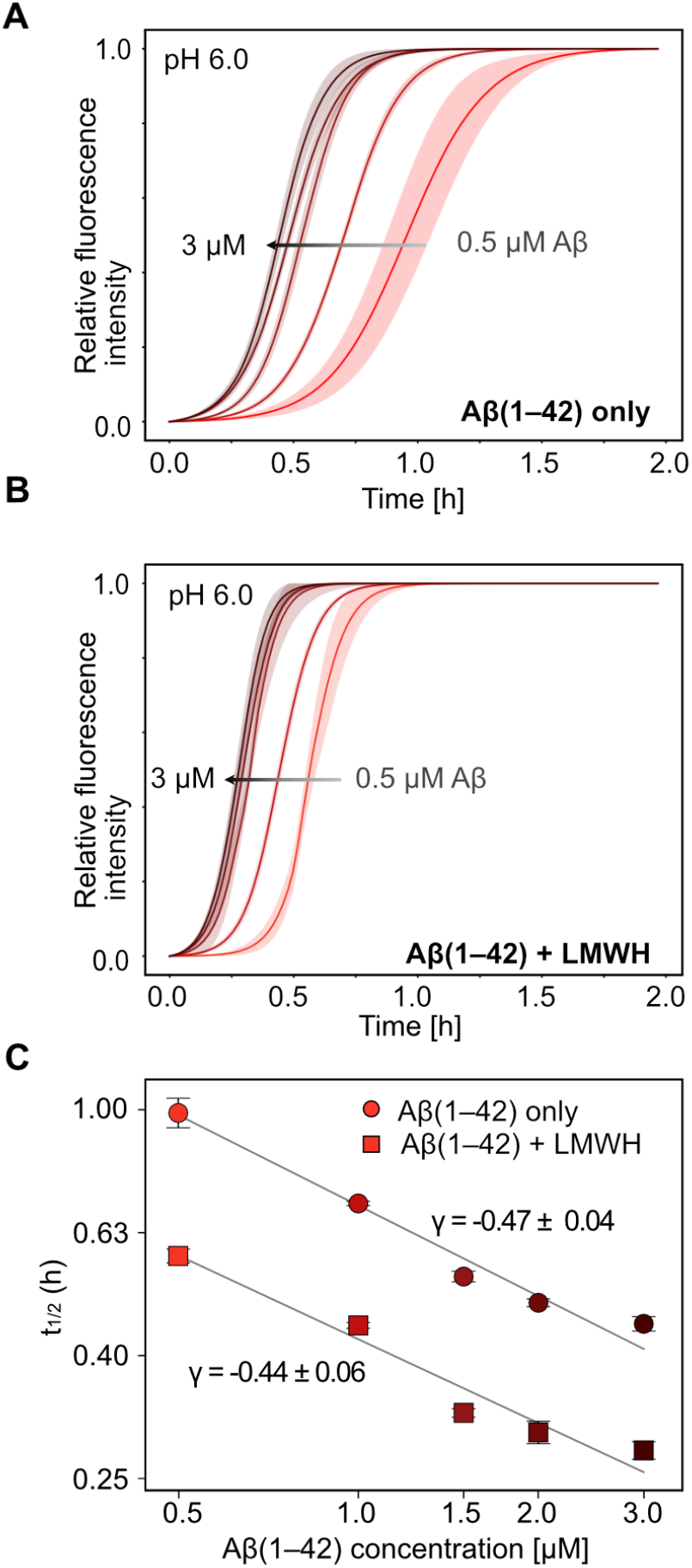
The effect of LMWH on the concentration dependence of amyloid aggregation kinetics. **A–B)** Amyloid formation kinetics at 0.5-3 µM of Aβ(1–42) in the absence (A) and presence (B) of LMWH. **C)** Aggregation half-times (t_*1/2*_) extracted from panels A and B, plotted against monomer Aβ(1–42) concentration (*m*_*0*_) in a log-log representation . The data series are fitted to a power law *t*_*1/2*_ = A *m*_*0*_ ^γ^.

Addition of LMWH does not significantly alter γ, indicating that the dominant aggregation mechanism remains unchanged. However, the absolute rates are systematically increased, as evidenced by a uniform shift of *t*_*1/2*_ towards lower values across all Aβ concentrations (**Figure 4C**). This acceleration is accompanied by a corresponding increase in the maximum aggregation rate *r*_*max*_ (**Figure S17**), consistent with the expected interdependence of *t*_*1/2*_ and *r*_*max*_ for aggregation dominated by secondary nucleation.^61^ Data obtained in the absence and presence of LMWH cluster separately but fall along the same trend line (**Figure S18**), further supporting a common underlying aggregation mechanism with and without LMWH. This shift in aggregation rate is consistent with the interpretation that the peptide experiences a higher effective concentration in the presence of the polyanionic GAG scaffold than in the bulk. In this interpretation, the shift in Figure 4C indicates that the effective concentration of Aβ in the presence of LMWH is approximately threefold higher than in the GAG-free control experiment (**Figure S19**). It should, however, be noted that the effect could equally stem from a decrease in the kinetic prefactor A, *i*.*e*. an increase in one or several underlying microscopic rate constants, which would be mathematically equivalent to an increase in effective *m*_*0*_. Such effects could, for example, be achieved by the GAG facilitating folding of the peptide into a more nucleation-competent conformation, as also suggested by MD simulations. The effect observed in the ensemble measurements is hence probably a combination of the two effects.

### Aggregation enhancement depends on multivalent peptide–GAG interactions

Key properties of GAGs arise from their polyelectrolyte nature, which enables multivalent interactions with proteins.^5^ Variations in chain length and degree of sulfation, therefore, provide opportunities for different modes of interactions with Aβ. In particular, the proposed mechanism in which GAGs promote peptide clustering implies a strong dependence on polymer length.

To investigate these effects systematically, we applied multidimensional separation to fractionate LMWH into well-defined populations based on size and net charge. Size-exclusion chromatography (SEC) first resolved the sample into GAG chains spanning 4 to 16 monosaccharide units (dp4–dp16) (**Figure 5A**). The fractions could be reliably assigned up to dp14, whereas the next fraction likely contains a mixture of longer chains ≥dp16. Equimolar amounts of the isolated GAG fractions were then incubated together with Aβ(1–42) and analyzed by the ThT aggregation assay. A clear dependence on GAG chain length was observed, with progressively stronger acceleration of aggregation for longer chain lengths (**Figure 5B**). A systematic decrease was evident as polymer length increased, yielding an approximately 40% reduction in the aggregation halftime between dp4 and dp16 (**Figure S20A**). Consistently, *r*_*max*_ increased steeply as GAG length increased (**Figure S20B**). For both parameters, an apparent saturation of the effect could be observed for the longest chain lengths.

**Figure 5.**
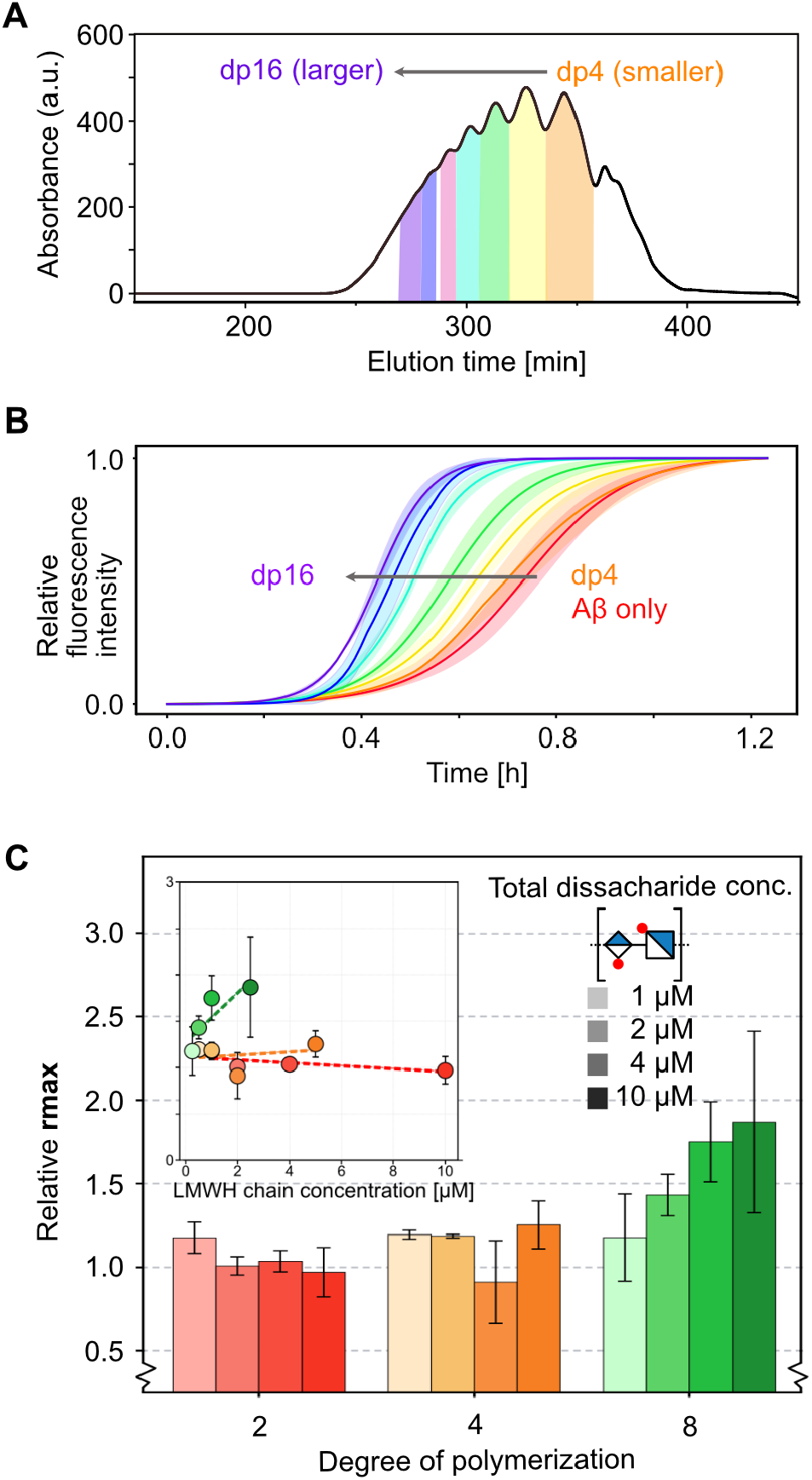
The effect of GAG chain length on Aβ aggregation. **A)** Size fractionation of LMWH by size exclusion chromatography into fractions of distinct chain length (degree of polymerization, dp), indicated by the colors: dp4 (orange), dp6 (yellow), dp8 (green), dp10 (light blue), dp12 (pink), dp14 (dark blue), and ≥dp16 (purple). **B)** ThT aggregation kinetics of 1 µM Aβ(1–42), without addition of heparin (red) and after equimolar additions of distinct chain lengths of LMWH (orange–purple) according to the fractionation in panel A. **C)** Relative *r*_*max*_ comparison from different dp of HS and LMWH (HS dp2, LMWH dp4, and dp8) upon increase of the total disaccharide unit concentration (1-10 µM). The inset shows the corresponding *r*_*max*_ as a function of chain concentration, with the chain concentration being the disaccharide concentration / (degree of polymerization/2).

It could be argued that incubating Aβ with equimolar concentrations of GAG chains of increasing length results in a higher total number of monosaccharide units in samples containing longer chains. To distinguish between an effect arising from total GAG content and one intrinsic to chain length, we next compared the fractionated LMWH dp4 and dp8 fractions, as well as the free HS disaccharide (dp2, the smallest unit in a GAG), while varying the total disaccharide unit concentration for each chain length between 1 and 10 µM.

Plotting *r*_*max*_ as a function of increasing total disaccharide concentration revealed very weak concentration dependences for the dp2 and dp4 samples, with minimal changes in aggregation behavior across the tested concentration range of 1 and 10 µM disaccharide units (**Figure 5C**, red and orange). In contrast, dp8 exhibited a clear concentration-dependent enhancement of aggregation (**Figure 5C**, green). Notably, this effect occurs despite dp8 being present at lower molar chain concentrations compared to dp2 and dp4 at equivalent disaccharide levels (**Figure 5C**, inset). For example, 2.5 µM dp8 (equivalent to 10 µM disaccharide units) produced an approximately twofold increase in aggregation rate, whereas 10 µM of the free disaccharides (dp2) showed no significant effects. These findings suggest that the major contribution to the enhancing effect on aggregation arises from chain-length-dependent multivalency rather than the total number of charged sugar units in solution, thereby ruling out a simple ionic strength effect. Moreover, they demonstrate that a minimum GAG chain length is required to effectively promote aggregation, consistent with a threshold for multivalent peptide recruitment.

Having established that aggregation enhancement depends on chain-length-dependent multivalency, we next considered how the strength of these interactions is influenced by GAG charge density. While longer chains provide a higher total negative charge, effective peptide accommodation could rely on cooperative contributions from chain length and charge distribution, rather than on a single driving factor. To disentangle these contributions, we specifically examined the role of sulfation, which modulates the local charge density along the polymer and thus represents an additional key parameter governing GAG–protein interactions. To examine this, the LMWH dp12 fraction isolated by SEC was further fractionated by anion-exchange chromatography (AXC). AXC of dp12 yielded multiple peaks separated according to net charge (**Figure 6A**). In comparison, AXC fractionation of dp4 resulted in a chromatogram shifted toward lower elution volumes, consistent with a lower net charge, and displayed a narrower distribution (**Figure S21**), reflecting the more limited number of possible sulfation patterns in shorter chains. For dp12, the most abundant species (**Figure 6A**, blue) and a more highly charged fraction (**Figure 6A**, purple) were isolated, and their effects on Aβ aggregation were assessed using the ThT assay. The results show that the more highly sulfated dp12 heparin accelerates aggregation more strongly, as evidenced by shorter *t*_*1/2*_ and increased *r*_*max*_ compared to the lower-sulfated fraction (**Figure 6B, Figure S22**), highlighting the importance of GAG charge, at a fixed chain length, in promoting amyloid formation.

**Figure 6.**
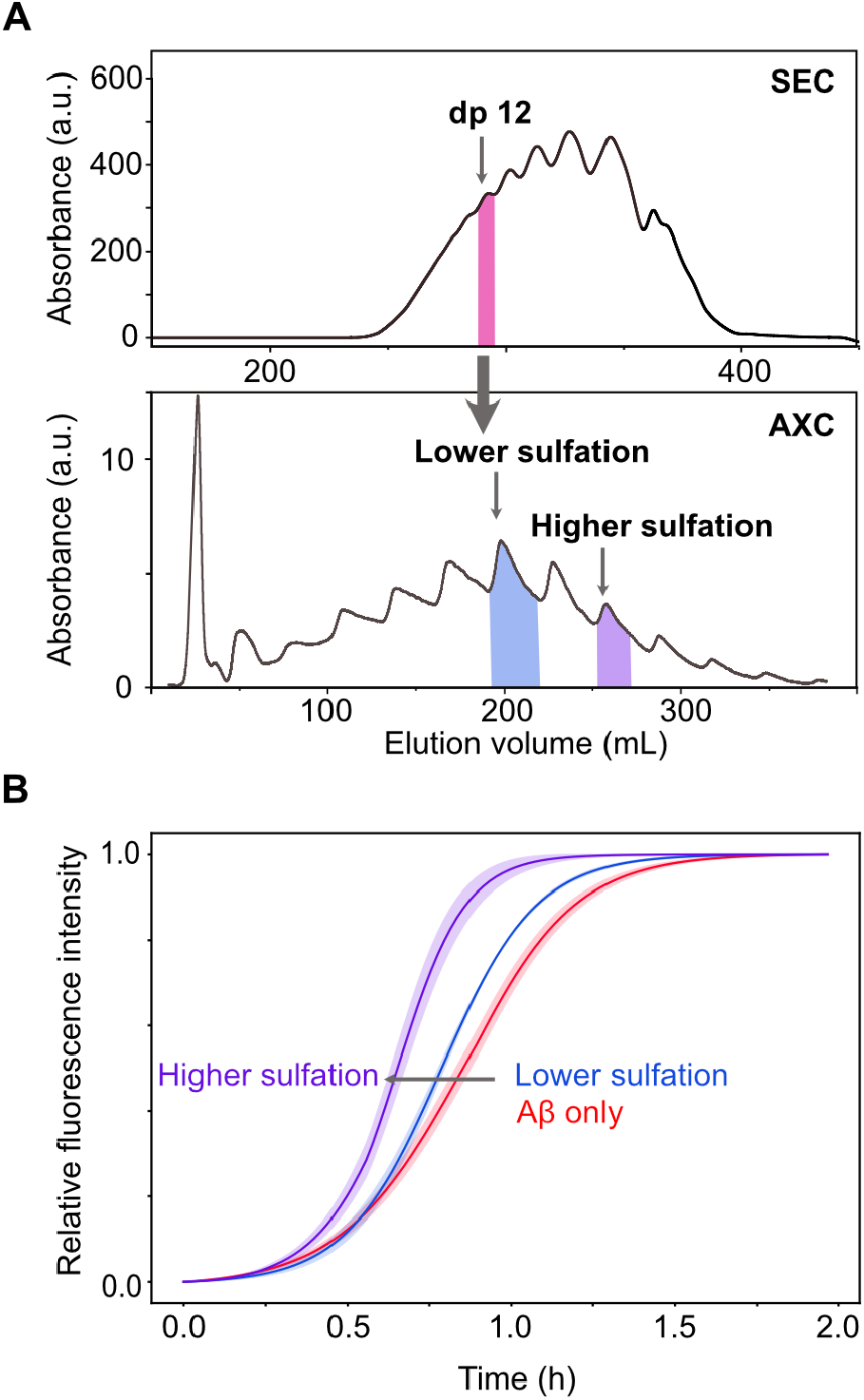
Effects of GAG sulfation on Aβ aggregation. **A)** Multidimensional separation of LMWH, first by size exclusion chromatography (SEC) followed by anion exchange chromatography (AXC) of the dp12 fraction from SEC into fractions of distinct degrees of sulfation. **B)** ThT assay of 1 µM Aβ(1–42) together with equimolar concentrations of dp12 heparin fractions with higher (purple) and lower (blue) degrees of sulfation.

Because other naturally occurring GAG classes are less highly sulfated than heparin, we next compared how different GAG types affect Aβ aggregation. Unsulfated hyaluronic acid (HA), as well as the more weakly sulfated chondroitin sulfate A (CSA) and dermatan sulfate (DS), were evaluated alongside highly sulfated heparin. All GAGs were size-fractionated by SEC, and dp12 fractions were used for the aggregation assays to ensure comparable chain length. A consistent trend was observed in which non- or weakly sulfated GAGs induced only modest reductions in *t*_*1/2*_ relative to Aβ alone, whereas highly sulfated heparin produced a substantially stronger acceleration of aggregation (**Figure S22**). The stronger aggregation-promoting effect of heparin compared with the other GAG types may reflect a combination of factors beyond overall sulfation, including glycan conformation, backbone rigidity, and sulfate-group presentation, for example the lower accessibility of 4-O-sulfation in CSA relative to 6-O-sulfation in heparin.

Together, these results demonstrate that GAG-mediated aggregation enhancement is governed by charge-dependent, multivalent interactions, with longer chains and higher sulfation promoting stronger effects by more efficiently clustering Aβ around the GAG scaffold.

## Discussion

We have shown here that GAGs are pH-dependent modulators of Aβ aggregation kinetics, without substantially altering the dominant aggregation mechanism or diverting aggregation away from β- sheet-rich amyloid fibril formation. Our results establish a mechanistic framework in which weak multivalent electrostatic interactions between GAGs and the pH-sensitive Aβ N-terminus promote peptide clustering facilitated by the hydrophobic C-terminal segment of the peptide. Such a switchable aggregation behavior provides an interesting mechanism for how the cell could be stressed during disease-related conditions, possibly triggering or enhancing the amyloid cascade observed in AD.

While both Aβ and GAGs are widely present in biological systems, aggregation may depend on local changes in cellular microenvironments. For example, alterations in GAG composition, including increased accumulation of heparan sulfate proteoglycans and altered sulfation patterns, have been associated with AD.^62,63^ In addition, altered trafficking of GAGs or Aβ may further contribute to this process,^64,65^ as Aβ is processed through endosomal–lysosomal pathways and disruptions in these trafficking routes are known to promote its accumulation and aggregation.^40,66^ Changes in extracellular pH, as observed during inflammation, could further contribute. Indeed, decreases in both brain tissue and cerebrospinal fluid pH have been observed in AD.^67^ This acidification is thought to be driven, at least in part, by metabolic dysfunction, as AD brains show elevated lactate levels, reflecting a shift toward glycolytic metabolism.^68^ In addition, activated microglia surrounding plaques contribute to the local microenvironment through inflammatory and metabolic activity, further promoting localized extracellular acidification.^69-71^

A central finding is the role of histidine protonation as a molecular switch that modulates electrostatic interactions with negatively charged sulfate groups on GAGs. Across GAGs with varying degrees of sulfation, our data indicate that sulfation strongly enhances GAG-mediated aggregation and likely strengthens binding, supporting a charge-driven interaction mechanism. These interactions occur *via* defined N-terminal motifs that resemble canonical heparin-binding sequences and are positioned directly adjacent to the central hydrophobic cluster (KLVFFA) in Aβ. This spatial proximity between the proposed GAG–recognition site and the aggregation hotspot likely facilitates coupling between GAG binding and peptide–peptide interactions. Importantly, our data show that GAGs do not fundamentally change the aggregation mechanism but instead increase the underlying rates. The net positive charge of the Aβ N-terminal segment and the net negative charge of the GAG chain arguably allow the GAGs to act as dynamic scaffolds that transiently concentrate multiple peptides within a confined volume, consistent with the apparent increase in effective peptide concentration. Such an aggregation environment would be important *in vivo*, where the available Aβ concentration is much lower than the concentrations needed for spontaneous amyloid formation in simple *in vitro* systems.^72^ It should be noted that the current study used free GAGs, whereas in biological systems, GAGs are also presented as proteoglycans at membrane surfaces. Membrane-anchored GAGs may further enhance local peptide concentration and further confine the interaction environments. In addition, membrane environments could upshift the pK_a_ of histidine residues, making protonation and hence peptide–GAG interactions yet more favorable.^53^

The dependence on GAG chain length and sulfation density further supports the proposed model. Short or weakly sulfated GAGs bind peptides but do not efficiently promote aggregation, whereas longer, highly sulfated chains enable multivalent recruitment and cooperative clustering. Together, these findings indicate that a threshold level of multivalency is required to transition from simple binding to aggregation-promoting behavior. The observed GAG binding is weak and occurs at a fast exchange rate on the NMR timescale, corresponding to ∼10^3^–10^5^ s^−1^ (μs–ms timescale).^73^ This likely gives rise to a rather “fuzzy” peptide coating of the GAG chain with constant exchange of bound and unbound species. Aggregation enhancement is therefore not proportional to binding affinity but depends on achieving sufficient multivalent peptide recruitment.

Very short GAGs, such as the disaccharide unit (dp2), are too small to facilitate peptide clustering. Molecular dynamics simulations of fondaparinux, comparable in size to dp4, indicate that this length primarily supports the binding of a single peptide, explaining its limited effect on aggregation compared to longer chains. Such longer GAG chains would, in contrast, enable simultaneous recruitment of multiple peptides, promoting cooperative assembly. Our data thus suggest a threshold chain length required for efficient aggregation enhancement, consistent with a transition from monovalent to multivalent binding. This observed threshold aligns closely with the intrinsic chain flexibility of the polymer, given that the persistence length of heparin is approximately 2.11 nm (roughly corresponding to dp4),^74^ oligosaccharides shorter than dp6 act as essentially rigid rods, limiting their structural capacity to recruit multiple peptide units. For highly sulfated variants, this persistence length can extend further, reinforcing a rigid conformation that restricts cooperative peptide recruitment at lower degrees of polymerization.^5^ Interestingly, the effect of chain length appears to plateau at higher degrees of polymerization, suggesting that beyond a certain length, additional increases in chain size no longer enhance clustering. This may reflect a balance between peptide recruitment and spatial dispersion along the polymer, particularly under conditions of limited peptide availability.

Our results help reconcile previously conflicting reports regarding the role of GAGs in Aβ aggregation.^9,44,45^ The present work suggests that GAG activity is highly context-dependent, with pH, multivalency, and GAG sulfation acting as key determinants of whether GAGs act as passive bystanders or active aggregation promoters. More broadly, our findings support a model in which GAGs act as dynamic scaffolds that concentrate and organize aggregation-prone peptides, thereby lowering the energetic barrier to amyloid formation, potentially contributing to the amyloid cascade during disease events. Our study further clarifies the role of the hydrophilic N-terminal segment as a modulating switch during aggregation. Notably, the identified GAG-binding region is not fully conserved across species: rodent Aβ contains substitutions within this segment (R5G, Y10F, H13R) that reduce its positive charge and alter its interaction properties, but does not contain any substitutions within the hydrophobic aggregation-prone regions.^75^ It may therefore be speculated that altered GAG-binding contributes to the reduced aggregation propensity and amyloid pathology observed in rodents.^76^ The GAG-binding motif further overlaps with well-known metal ion binding sites in the Aβ peptide,^77,78^ which are generally considered to slow down amyloid formation, suggesting a possibility of more complex interplay among peptide, GAGs, and other interaction partners.^21,78^ Importantly, this also suggests that disrupting electrostatic peptide–GAG interactions, such as through competitive binding to the N-terminal region of Aβ, may provide a strategy to attenuate GAG-triggered aggregation.

## Materials and Methods

### Glycosaminoglycan preparation

Enoxaparin sodium, Fondaparinux sodium, Dermatan sulfate, and Hyaluronic acid were purchased from Sigma Aldrich. Chondroitin sulfate A was purchased from Biosynth. Dp2 (ΔUA(2S)-GlcNAc(6S)) Heparan sulfate was purchased from Iduron (UK). CSA and DS were digested using Chondroitinase ABC (Sigma Aldrich) and HA was digested using hyaluronidase (Sigma Aldrich). For further details, see the extended materials and methods. Fractionation was performed on a Knauer Azura FPLC system (Berlin, Germany). GAGs were dissolved in SEC mobile phase (500 mM ammonium acetate) and were separated by SEC using a column packed with 350 mL of Superdex 30 resin (Cytiva, Germany).

Collected fractions were freeze-dried and redissolved in water to a concentration of 100 mM. Anion-exchange was performed using a column packed with the DEAE Sephacel weak anion resin (Cytiva, Germany). Samples were loaded in 50 mM Tris-base, pH 8, and eluted with a gradient from 250 mM to 2 M NaCl. Collected fractions were desalted using a Superose 6 column, concentrated by freeze-drying, and further purified by ethanol precipitation before being re-dissolved in water.

GAG concentrations were determined by their absorbance at 232 nm using a DS-11 microvolume spectrophotometer (DeNovix). A calibration curve was constructed using Chondroitin sulfate disaccharide (GlcA(2S)-GalNAc(4S)) (Iduron, UK).

### Peptide synthesis

Truncated and C-terminally amidated Aβ fragments Aβ(1–28) and Aβ(1–16), as well as histidine-to-alanine substitution variants (NoH) of these peptides, were synthesized by microwave-assisted Fmoc solid-phase peptide synthesis (SPPS) using a Liberty Blue 2.0 peptide synthesizer on Rink Amide ProTide® resin (0.2 mmol g^−1^ loading, 0.05 mmol scale). Amino acids were coupled using Oxyma/*N,N′*-diisopropylcarbodiimide (DIC), and Fmoc deprotection was performed with 20% piperidine in DMF containing 0.1 M Oxyma. After chain assembly, peptides were cleaved with a TFA-based cocktail, precipitated in cold diethyl ether, and dried under nitrogen. Crude peptides were purified by preparative reverse-phase HPLC and characterized by ESI-MS and analytical RP-HPLC.

### Peptide preparation

Recombinant Aβ(1–42) peptides were obtained from rPeptide (USA), and synthetic Aβ(12–28) was purchased from GenScript. Recombinant Aβ(1–42) and synthetic peptide fragments were dissolved in 8 M guanidinium chloride (pH 8.0) and subjected to size-exclusion chromatography (SEC) on Superdex 75 10/300 Increase or Superdex 30 10/300 Increase columns, equilibrated in 20 mM sodium phosphate buffer (pH 7.5), to isolate monomeric peptide fractions. Peptide concentrations were determined by an o-phthalaldehyde (OPA) fluorescence assay.^79^ SEC-filtered peptide aliquots were diluted to 80 µL and mixed with 5 µL of OPA solution (Thermo Scientific, USA) in a black 96-well half-area PEG-ylated low-binding polystyrene plate with transparent bottom (Corning, USA). Fluorescence was measured from the top (λ_ex_ = 340 nm, λ_em_ = 440 nm). Concentrations were determined using an Aβ(1–40) standard curve. Aliquots were kept on ice until further analysis.

### Thioflavin T aggregation assay

Thioflavin T (ThT) aggregation assays were performed in 96-well half-area PEG-ylated low-binding polystyrene plates, black with transparent bottom (Corning, USA). Freshly SEC-purified Aβ(1–42) was further diluted in 20 mM sodium phosphate at pH 6.0 or pH 7.4, as indicated, and supplemented with 80 µM ThT (abcr Chem) before loading 50 µL into each well of the microwell plate. Samples were aggregated in triplicate under quiescent conditions at 37°C in a Spark multimode plate reader (Tecan, Switzerland). ThT was excited at 440 nm, and its fluorescence emission was monitored every minute at 480 nm from the bottom of the plate.

The triplicate data were fitted to a sigmoidal model to obtain the aggregation half-time (*t*_*1/2*_):

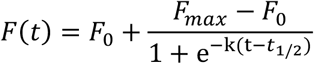

The *r*_*max*_ is calculated by taking the first derivative of F(t) where for the normalized data, the *r*_*max*_ is equal to k/4:

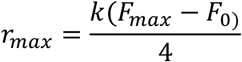

### Cryo-electron microscopy

Cryo-transmission electron microscopy (Cryo-TEM) was performed using 20 µM Aβ(1–42) with or without 20 µM LMWH in a 20 mM sodium phosphate buffer, pH 6, and the sample was incubated for 2 days at 37°C. Perforated carbon film-covered microscopical grids (200 mesh, R1.2/1.3 batch of Quantifoil, MicroTools GmbH, Jena, Germany) were cleaned with chloroform and hydrophilized by 60 s glow discharging at 8 W in a BALTEC MED 020 device (Leica Microsystems, Wetzlar, Germany) before use. 3.8 μL aliquots of the sample solutions were applied to these grids, and automatically blotted and plunge frozen in a Vitrobot Mark IV (Thermo Fisher Scientific Inc., Waltham, Massachusetts, USA) using liquid ethane as cryogen. The frozen grids were assembled into autogrids and transferred to a Talos Arctica electron microscope equipped with a high-brightness field-emission gun (XFEG) operated at 200 kV acceleration voltage. Micrographs were recorded on a Falcon 3 direct electron detector (Thermo Fisher Scientific Inc., Waltham, Massachusetts, USA) at a nominal magnification of 28,000× (0.375 nm/dot).

### Surface Plasmon Resonance

Interaction studies were performed on a Biacore X100 (GE Healthcare) at 25°C. A CM5 sensor chip was activated using standard EDC/NHS chemistry to immobilize streptavidin (100 µg/mL in 10 mM NaOAc, pH 5.0), then quenched with 1 M ethanolamine. Biotinylated Enoxaparin (dp14) was subsequently captured on the streptavidin surface (50 RU). For kinetic analysis, Aβ(1–16) was injected in duplicate at a flow rate of 10 µL/min (180 s association, 260 s dissociation) in 20 mM sodium phosphate buffer at various pH values. The surface was regenerated between cycles with 30 s pulses of 2 M NaCl in 50 mM NaOH. The binding constant K_D_ was analyzed using Tracedrawer and fitted to a 1:1 binding model.

### Circular Dichroism spectroscopy

CD spectra of Aβ(1–28) and Aβ(1–28) NoH were recorded on a Jasco J-810 spectropolarimeter equipped with a Jasco PTC-432S Peltier temperature control system. All measurements were conducted at 37 °C. CD spectra of Aβ(1–42) with the addition of LMWH were measured with an Olis DSM20 Spectrophotometer. Each spectrum was background-corrected using a subtraction of the buffer spectra measured under the same conditions, and the mean of three consecutive measurements was taken. For Aβ(1–28), low concentrations of 15 µM were measured using a 2 mm cuvette, concentrations of 30 µM were measured using a 1 mm cuvette, and high concentrations of 100 µM were measured with a demountable 0.5 mm cuvette. 20 µM of Aβ(1–42) was measured using a 1 mm cuvette. Each spectrum was normalized by path-length, molar concentration, and the number of amide bonds. Each Aβ(1–28) sample was prepared by dissolving the peptide in a sodium phosphate buffer for pH-dependent measurements. The Aβ(1–42) sample was prepared by diluting the stock solution from the SEC in 20 mM sodium phosphate buffer (pH 6) to the desired concentration. Samples were sonicated before the first measurement and incubated at 37 °C between measurements.

### NMR spectroscopy

NMR spectra were recorded on a Bruker Avance III 700 MHz spectrometer equipped with a cryoprobe at 278 K in 20 mM sodium phosphate buffer (pH 6.0, 10% D_2_O). Backbone and selected side-chain resonances of Aβ(1–28) were assigned using standard 2D experiments (NOESY, TOCSY, HSQC, HMQC) in combination with literature data and predicted chemical shifts.^55 80^ Peptide–GAG interactions were probed by titrating fondaparinux or enoxaparin into peptide samples and monitoring chemical shift perturbations. Binding affinities were estimated by fitting concentration-dependent chemical shift changes to a one-site binding model. Spectra were processed with TopSpin 3.2 (Bruker, Billerica, USA) and analyzed with CcpNmr Analysis version 3 and MestReNova version 14.0.3-30573.^81^

### Molecular Dynamics simulations

Initial extended structures of Aβ(1–28) and its sequence variants were generated in PyMOL, and histidine protonation states corresponding to pH 6 were assigned using H++.^82^ The C-terminus was amidated while the N-terminus retained its protonated amine. Fondaparinux coordinates were obtained from the RCSB database and parameterized using the Glycan Reader & Modeler module of CHARMM-GUI.^83^ The resulting parameters for fondaparinux were validated as described in the SI. Simulations were performed in GROMACS^84^ with the CHARMM36 force field.^85^ Systems were solvated in explicit TIP3P water within a dodecahedral box^86^ and neutralized with Na^+^/Cl^−^ ions, with electrostatics treated using the particle-mesh Ewald method under periodic boundary conditions.^87^ After steepest-descent energy minimization, systems were equilibrated at 300 K and 1 bar in the NPT ensemble using velocity-rescaling thermostatting^88^ and a Parrinello–Rahman barostat,^89^ followed by production runs employing a Nosé–Hoover thermostat.^90^ Three independent 1-µs trajectories were generated for each peptide (monomers or trimers) in the absence and presence of fondaparinux, with coordinates saved every 20 ps for analysis. Residue–residue contact analysis was performed using the MDAnalysis library. A contact was defined when the minimum interatomic distance between a residue pair fell below a 1.0 nm threshold. The contact occupancy was subsequently determined as the fraction of trajectory frames satisfying this criterion, yielding a time-averaged contact map. The conformational preferences of Aβ(1–28) were characterized by transition networks using the ATRANET framework.^91^ It computes transition matrices from MD trajectories using descriptors, for which we used the end-to-end distance and numbers of α-helical and β-sheet residues^92^ calculated with DSSP.^93^ Networks were visualized and analyzed in Gephi.^94^

## Supporting information

Extended methodology and Supplementary Figures

## Acknowledgements

This work was supported by DFG, the German Research Foundation, through GRK2662, grant number 434130070 to EM, CR, CK, BK, and KP. YW and BStr gratefully acknowledge computing time on the supercomputer JURECA at Forschungszentrum Jülich under grant AMYAI. YW is supported by a PhD fellowship from the Chinese Scholarship Council. NÖ is supported by a fellowship from the Wenner-Gren Foundations. We acknowledge the research infrastructure and support provided by the SupraFAB Research Facility, funded by the German Federal Government and the State of Berlin. We further acknowledge the assistance of the NMR Core Facility at BioSupraMol, Freie Universität Berlin, supported by the DFG (the German Research Foundation). Dr. Mario Schubert is thanked for the valuable discussion regarding the NMR analysis.

## Author contributions

NÖ and KP conceived and designed the project. EM, CR, JS, FF, CK, WP, BSch, AP, and RZ performed experiments. YW and SS performed simulations. EM, CR, JS, YW, GPS, and NÖ analyzed and visualized data. GMV, BStr, BK, KP, and NÖ supervised the work. BStr, BK, and KP provided resources and funding for the project. NÖ coordinated the project. EM and NÖ wrote the manuscript with input from all other authors.

