## Extended methodology and Supplementary Figures for "Glycosaminoglycans Promote Amyloid-β Aggregation via Multivalent, pH-Dependent Interactions"

### A. Extended Materials and Methods

#### A.1. Chemical synthesis of Aβ peptide fragments

##### A.1.1 Synthesis

Truncated and C-terminally amidated Aβ fragments Aβ(1–28) and Aβ(1–16), as well as histidine-to-alanine substitution variants, were synthesized by microwave-assisted Fmoc SPPS using a Liberty Blue 2.0 peptide synthesizer. Synthesis was carried out on Rink Amide ProTide® resin (loading 0.2 mmol $g^{-1}$) at a 0.05 mmol scale from the C- to the N-terminus following standard Fmoc chemistry. Amino acids were activated with Oxyma and *N,N′*-diisopropylcarbodiimide (DIC), and Fmoc deprotection employed 20% piperidine in DMF containing 0.1 M Oxyma to enhance deprotection efficiency.

Microwave-assisted conditions were optimized to reduce on-resin aggregation and improve coupling efficiency. Sequence-dependent adjustments—including prolonged or double coupling—were introduced for residues prone to incomplete coupling or side reactions, such as histidine, aspartate-containing motifs susceptible to aspartimide formation, sterically hindered arginine, and the N-terminal residue.

Following chain assembly, peptides were cleaved from the resin using a TFA-based cocktail (94% TFA, 2.5% ethanedithiol, 2.5% water, 1% triisopropylsilane) for 2 h at room temperature. The cleavage mixture was precipitated with cold diethyl ether, centrifuged, washed repeatedly with ether, and dried under a gentle nitrogen stream to yield crude peptide.

##### A.1.2 Purification and analytical characterization

Purification of crude peptides was performed on a Nexera Prep preparative HPLC system equipped with UV detection (10 mm flow cell) and automated liquid handling. Dissolved peptide samples were filtered through a 0.45 µm PTFE membrane prior to injection. Separation was achieved on a Kinetex® C18 column (5 µm, 100 Å, 250 × 21.2 mm) using water (eluent A) and acetonitrile (eluent B), each containing 0.1% TFA, at a flow rate of 10 mL $min^{-1}$. Peptides were purified using a linear gradient from 10% to 45% B over 25 min, monitored at 220 nm. Fractions were collected, analyzed by analytical RP-HPLC, and pure fractions were combined. Acetonitrile was removed under reduced pressure, and aqueous peptide solutions were frozen in liquid nitrogen and lyophilized.

Analytical RP-HPLC was performed on a VWR-Hitachi Chromaster system using a Kinetex® C18 column (250 × 4.6 mm, 5 µm, 100 Å) with water/acetonitrile (0.1% TFA) and a 5–50% B gradient over 18 min at 1 mL $min^{-1}$, detected at 220 nm. Peptide identity was confirmed by electrospray ionization Time-of-Flight (ESI-TOF) mass spectrometry on an Agilent 6320 System after being dissolved in 10% HFIP/90% water.

Purified peptides were dissolved in hexafluoroisopropanol (HFIP), sonicated, filtered, and dried prior to redissolution in guanidinium-containing phosphate buffer. Concentrations were determined by UV absorbance.

### A.2 Enzymatic digestion of different GAGs

100 mg of CSA dissolved in 7.6 mL of water, and 1 mL of 100 mM sodium phosphate (Sigma Aldrich) and 750 uL of 2M NaCl were added. 0.5 U of Chondroitinase ABC (Sigma Aldrich) was dissolved in PBS to a final volume of 625 µL and added to the mixture. After an hour incubation at 37 °C, the mixture was immediately freeze-dried. The same depolymerization was performed for DS. 10 mg of HA was dissolved in 10 mL of 100 mM Ammonium acetate, pH 4 buffer with 150 mM NaCl. Hyaluronidase, purchased from Sigma Aldrich was added in a 1:80 (enzyme: substrate) ratio. The reaction was left at 37 °C for 24 hours and immediately freeze-dried.

#### A.3 Surface Plasmon Resonance

Enoxaparin dp 14 was biotinylated with biotin aminooxy (Sigma Aldrich) during incubation in 10 mM NaOAc, pH 5.5 buffer for 5 days. The biotinylated Enoxaparin was purified by SEC and lyophilized. The CM5 biosensor chip from GE Healthcare (Germany) was activated with 1-Ethyl-3-(3-dimethylaminopropyl)carbodiimide (EDC) and N-hydroxysuccinimide (NHS) (GE Healthcare, Germany) and 130 µL solution of Streptavidin (100 µg/mL in 10 mM NaOAc, pH 5) was injected at a flow rate of 5 µL/min. The remaining active ester was quenched with 1 M Ethanolamine solution. Then, 130 µL biotinylated Enoxaparin in HBS-EP buffer (150 mM NaCl, 10 mM HEPES, 3 mM EDTA, 0.05% P20, pH 7.4). was injected onto the streptavidin-modified chip, which resulted in 50 RU. The chemicals NaCl, HEPES, and EDTA were purchased from Carl Roth.

The experiment was performed in a 20 mM Sodium phosphate buffer at different pHs. The buffer was prepared with Sodium dihydrogen phosphate monohydrate and Sodium phosphate dibasic dihydrate purchased from Sigma Aldrich. 2 M NaCl in 50 mM NaOH solution was used in 30s pulses for regeneration at the end of each cycle.

#### A.4 NMR spectroscopy

NMR spectra were acquired on a Bruker Avance III 700 MHz spectrometer equipped with a 5 mm triple resonance cryoprobe. The NMR measurements were performed at 278 K in a 20 mM sodium phosphate buffer, pH 6.0, with 10% $D_2O$. Fondaparinux and Enoxaparin were prepared in the same buffer. As histidines are highly sensitive to pH changes, the pH of the NMR samples was controlled.

Assignments of backbone amide as well as selected side chain proton resonances were obtained by using $^1H$-$^1H$-NOESY (80 ms mixing time), $^1H$-$^1H$-TOCSY (MLEV-16 120 ms mixing time), $^1H$-$^{15}N$-HSQC, and $^1H$-$^{13}C$-HMQC spectra of 100 μM unlabeled Aβ(1-28), also taking into account previously published chemical-shift assignments for Aβ(1-40) at pH7.4 and 278 K[1] and Aβ (1-42) at pH 7.4 and 273 K (BMRB 25218), as well as chemical shifts predicted for Aβ(1-28) by ncIDP.[2] A $^1H$-$^{15}N$-HSQC spectrum was also measured of 100 μM Aβ(1–28) in the presence of 1 mM fondaparinux.

Titrations were performed by adding highly concentrated GAGs to Aβ-alone NMR samples, resulting in 33 μM Aβ(1-28) with 0 μM, 33 μM, 66 μM, 165 μM, 330 μM, 1000 μM, 1542 μM Fondaparinux; 50 μM Aβ(1-28) with 25 μM, 50 μM, 100 μM, 250 μM, 1000 μM Enoxaparin; 50 μM NoH Aβ(1-28) with 0 μM, 50 μM, 250 μM Fondaparinux; 50 μM Aβ(1-16) with 0 μM, 25 μM, 50 μM, 100 μM, 250 μM, 1000 μM Enoxaparin; and 25 μM Aβ(12-28) with 12.5 μM, 25 μM, 50 μM, 125 μM, 500 μM Enoxaparin.

Chemical shift perturbation was calculated according to equations shown below for one-dimensional proton spectra:

$$\Delta\delta(^1H) = \sqrt{(\Delta\delta(^1H))^2}$$

and for $^1H$-$^{15}N$-HSQC spectra:

$$\Delta\delta(^1H, ^{15}N) = \sqrt{(\Delta\delta(^1H))^2 + (0.15 \cdot \Delta\delta(^{15}N))^2}$$

$K_D$ values were estimated based on fitting the chemical shift distance in dependence of GAG concentration to a one-site binding model using OriginPro 2010b.

### A.5 Molecular dynamics simulations

#### A.5.1 System preparation and simulation protocol

Initial extended structures of wild-type Aβ(1–28) and Aβ(1–28) H6A/H13A/H14A (NoH) were generated in PyMOL. Histidine protonation states corresponding to pH 6 were assigned using the H++ web server,[3] and the C-terminus was amidated while the N-terminus retained its protonated amine. Fondaparinux coordinates were obtained from the RCSB database and parameterized using the Glycan Reader & Modeler module of CHARMM-GUI.[4]

Simulations were carried out in GROMACS[5] using the CHARMM36 force field.[6] Each system was placed in a dodecahedral box (~1310 $nm^3$), solvated with explicit TIP3P water,[7] and neutralized with $Na^+$ and $Cl^-$ ions (~0.02 M). Long-range electrostatics were treated using the particle-mesh Ewald method under periodic boundary conditions.[8] Energy minimization employed the steepest-descent algorithm, followed by equilibration at 300 K and 1 bar in the NPT ensemble using a velocity-rescaling thermostat[9] and Parrinello–Rahman barostat.[10] Production simulations used the Nosé–Hoover thermostat[11] together with the Parrinello–Rahman barostat.

For monomer simulations, three independent 1-µs trajectories were generated for wild-type Aβ(1–28) and NoH in the absence and presence of fondaparinux. For multipeptide simulations, three independent 1-µs trajectories were generated for systems containing three Aβ(1–28) peptides in the absence or presence of one fondaparinux molecule. A 500-ns fondaparinux-only simulation was performed for force-field validation. Coordinates were saved every 20 ps for analysis, yielding 25,000 frames for the 500-ns simulation and 50,000 frames for each 1-µs trajectory.

#### A.5.2 Validation of fondaparinux force-field parameters

Force-field accuracy was assessed by comparison with published structural data through calculation of glycosidic dihedral angle distributions, Cremer–Pople ring-puckering coordinates, and three-bond J-couplings derived from Karplus relationships. These quantities were computed using AmberTools cpptraj analysis routines.

#### A.5.3 Contact maps

Intermolecular peptide–GAG and peptide–peptide contacts were analyzed from the MD trajectories using distance-based contact analysis implemented in Python with MDAnalysis. For peptide–GAG contact maps, contacts were calculated between peptide residues and individual fondaparinux monosaccharide units. For peptide–peptide contact maps, contacts were calculated between residue pairs belonging to different peptide chains.

A contact was assigned in a given trajectory frame when the minimum interatomic distance between the two analyzed groups was below 1.0 nm. Contact occupancies were then calculated as the fraction of trajectory frames in which this criterion was satisfied. The resulting occupancies were used to generate time-averaged contact maps and one-dimensional contact-frequency profiles. Contact frequencies were averaged over independent trajectories unless otherwise stated. Absolute or normalized contact frequencies are reported as indicated in the corresponding figure captions.

#### A.5.4 Transition-networks

Conformational transition networks were generated using the ATRANET framework, which computes transition matrices from MD trajectories using descriptors including end-to-end distance (dNC) and numbers of α-helical (Nα) and β-sheet (Nβ) residues. Networks were visualized and analyzed in Gephi using Yifan Hu and ForceAtlas2 layout algorithms, with node size scaled by population and modularity analysis applied to identify structural clusters.

### B. Supporting Figures

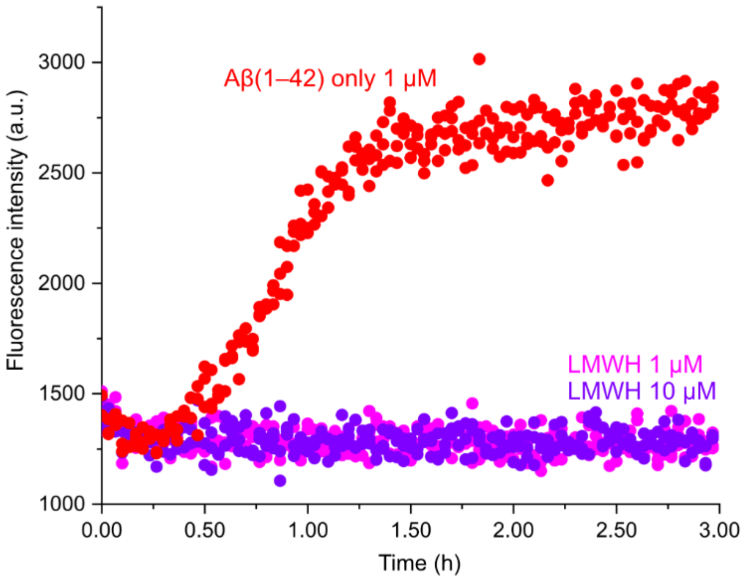

**Figure S1.** LMWH does not interact with Thioflavin T, performed at pH 6.0. Aβ(1–42) ThT data is shown as a comparison.

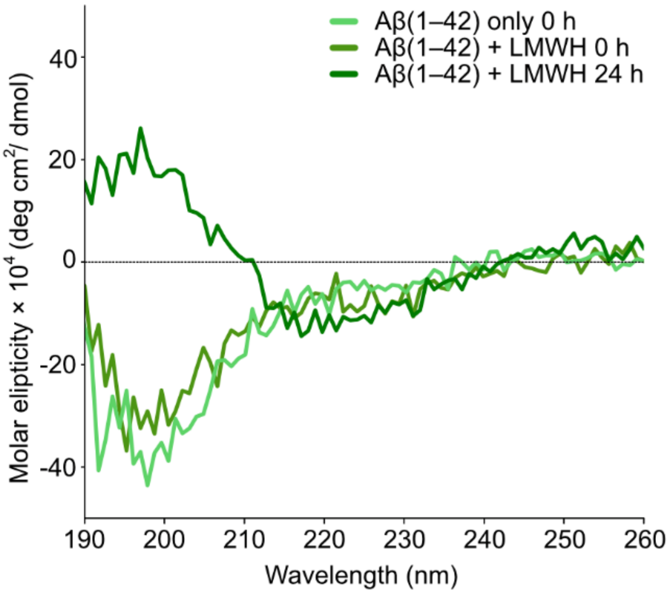

**Figure S2**: CD spectra of Aβ(1–42) (20 μM) incubated with LMWH (20 μM) at pH 6 for 24 hrs.

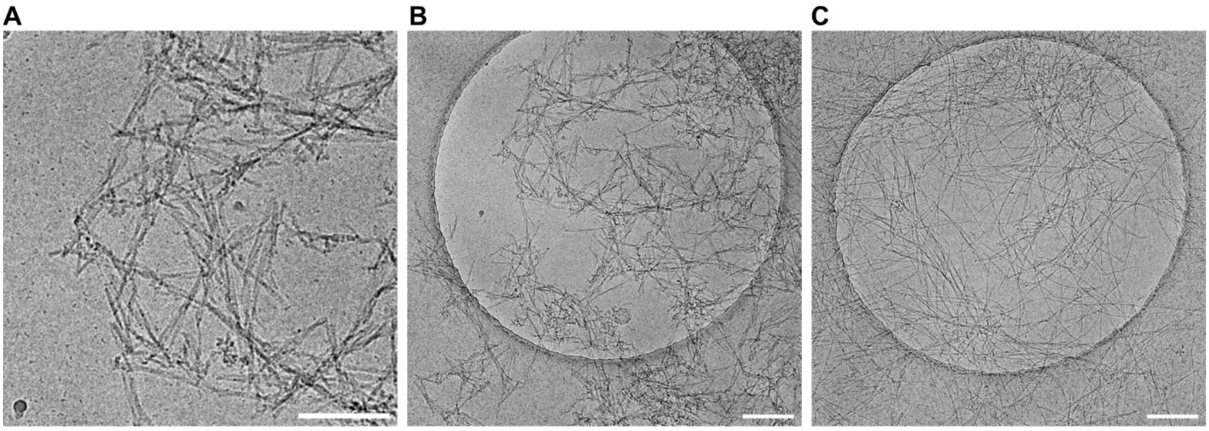

**Figure S3**. Cryo-EM micrograph of **(A)** and **(B)** Aβ(1–42) only, and **(C)** Aβ(1–42) with LMWH. Scale bars: 50 nm **(A)**, 200 nm **(B, C)**

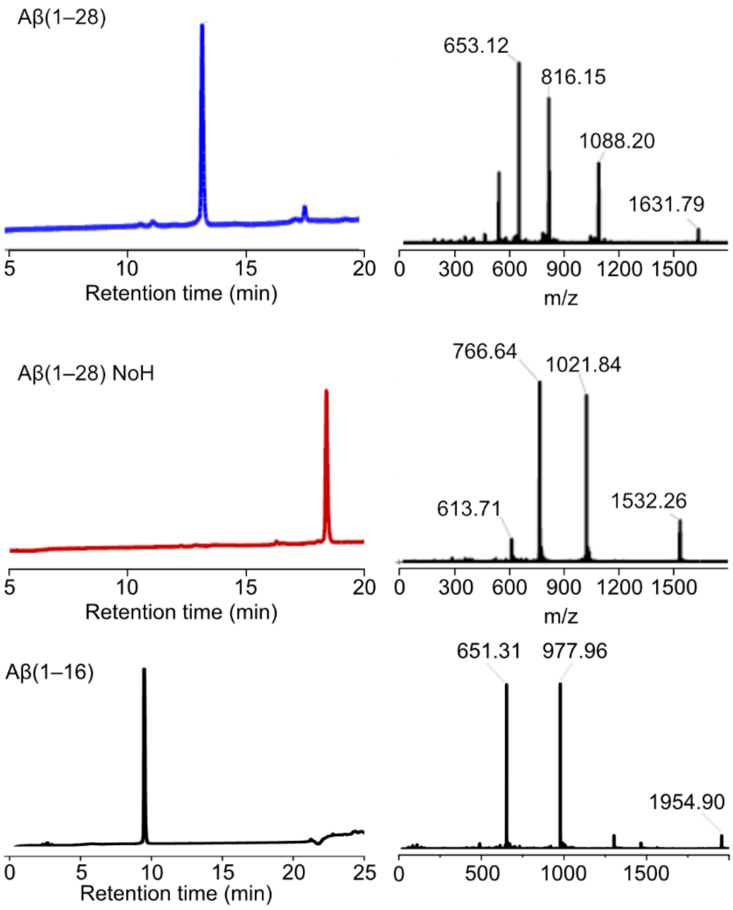

**Figure S4.** Analytical HPLC chromatograms (l.) and ESI-TOF mass spectra (r.) of purified peptides. Peptides were purified with a linear gradient from 10–45 % B over 25 mins, 1.00 mL/min.

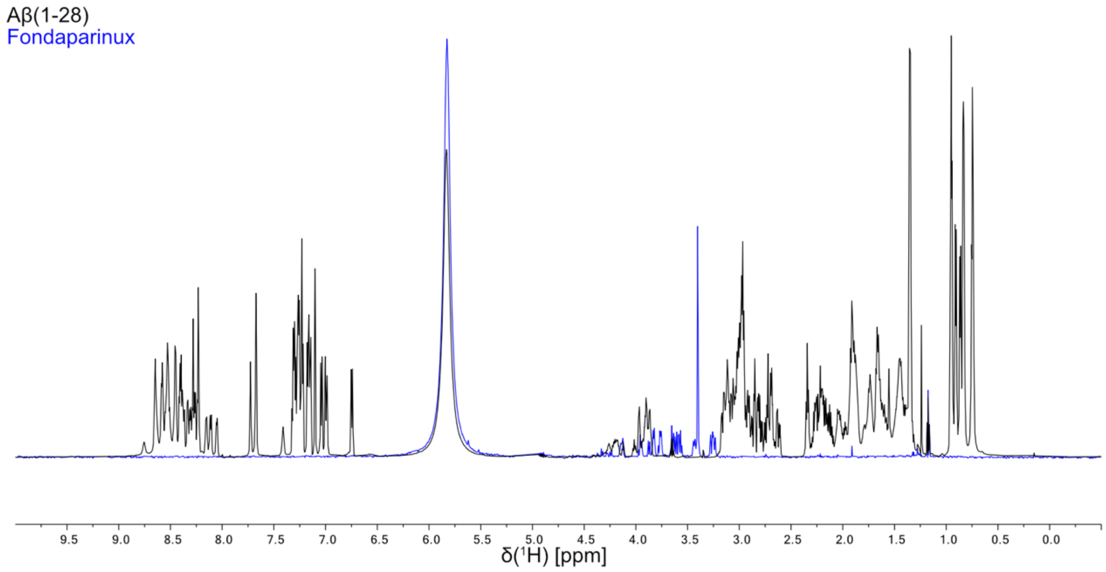

**Figure S5.** $^1$H NMR spectra of 50 μM Fondaparinux (blue) and 50 μM Aβ(1–28) (black) in 20 mM sodium phosphate buffer pH 6.0 shown as overlay. The characteristic regions of Fondaparinux and Aβ(1–28) do not overlap.

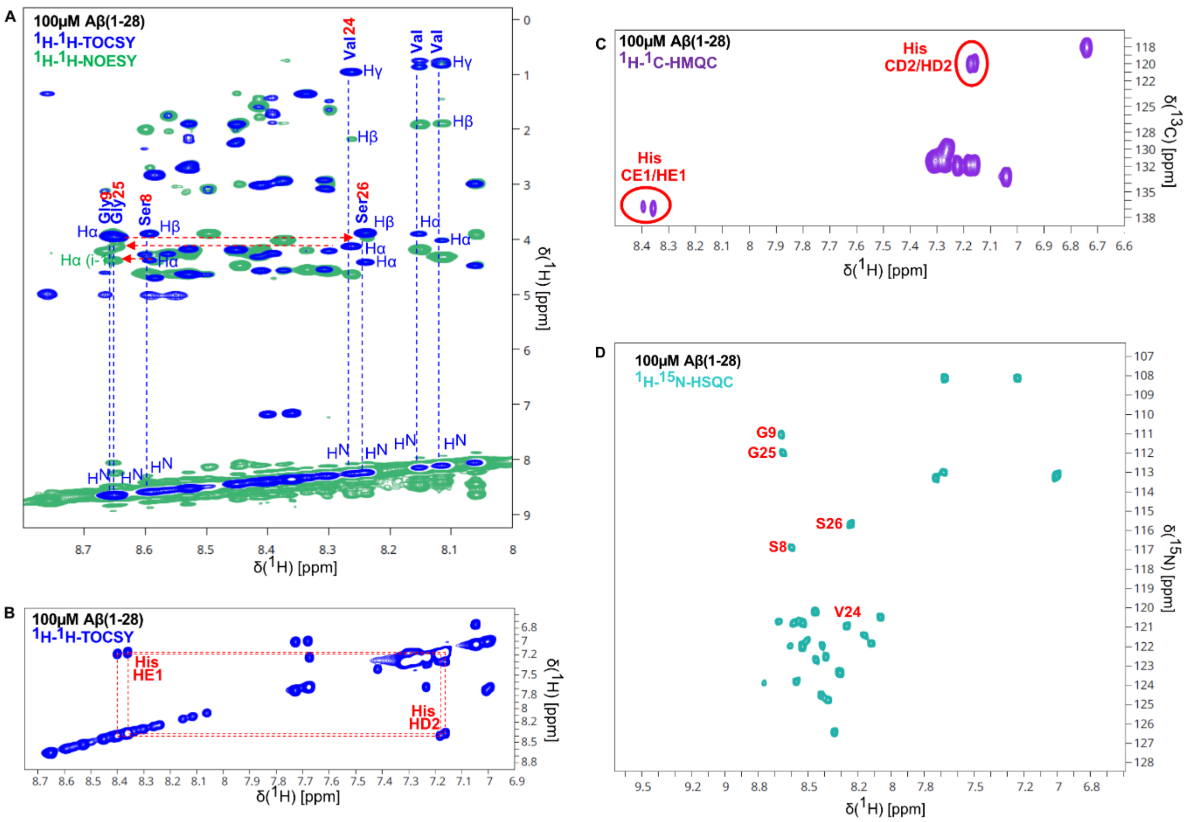

**Figure S6.** Assignment of backbone amide proton and selected side chain resonances of 100 µM Aβ(1–28) at pH 6. **(A)** Assignment procedure using sequential NH-Hα cross peaks exemplified for some residues in $^{1}$H-$^{1}$H-NOESY and $^{1}$H-$^{1}$H-TOCSY spectra. B,C) Identification of Histidine HE1 and HD2 side chain signals based on $^{1}$H-$^{1}$H-TOCSY **(B)** and $^{1}$H-$^{13}$C-HMQC **(C)** spectra. **(D)** Assignments exemplified in A) shown in $^{1}$H-$^{15}$N-HSQC spectrum.

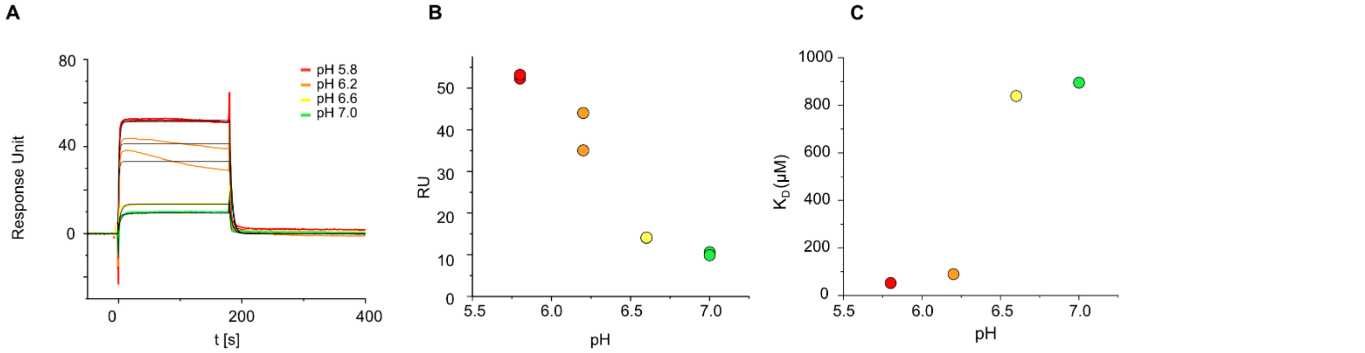

**Figure S7.** SPR pH titration across pH 5.8 – 7.0 with Aβ(1–16) on immobilized LMWH, shown as **(A)** raw data **(B)** comparison of maximum response unit against pH, and **(C)** the fitted $K_D$ from duplicates against pH.

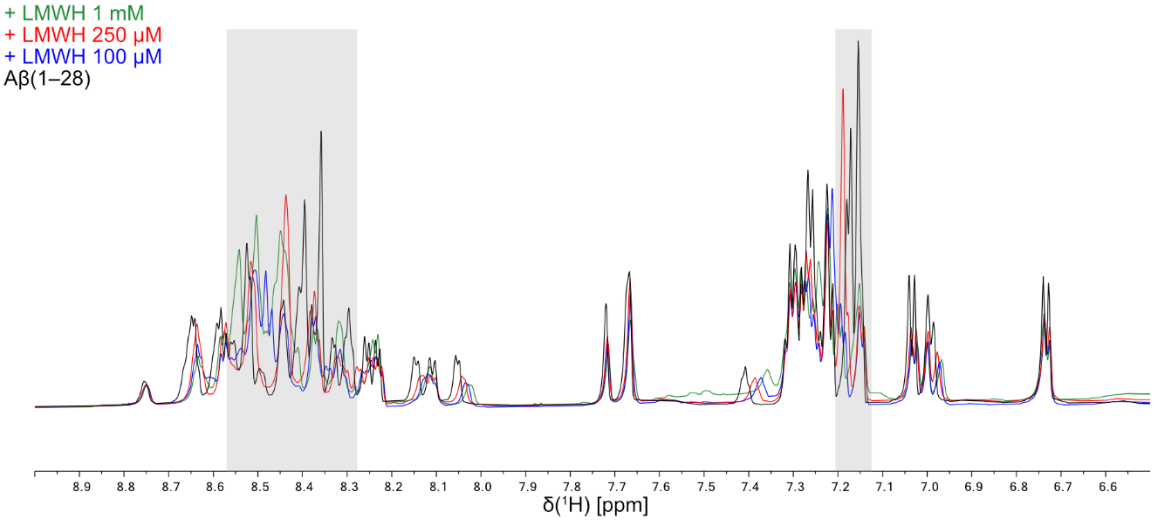

**Figure S8.** $^{1}$H NMR intensity of 50 µM Aβ(1–28) decreases with increasing LMWH concentration. The grey highlighted area is the chemical shift of Histidine HE1 and HD2 protons. Spectra were recorded in 20 mM sodium phosphate buffer pH 6.0.

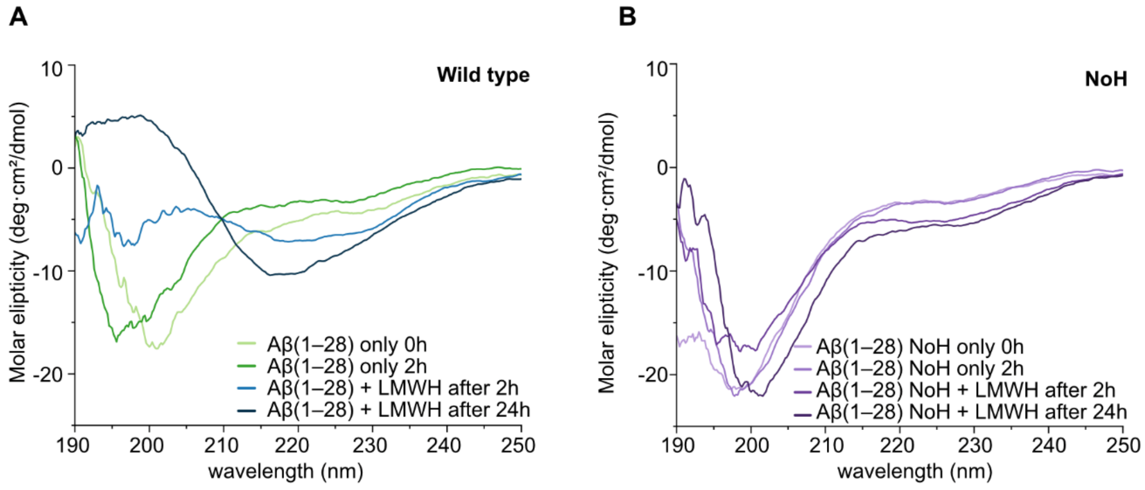

**Figure S9** CD spectra of **(A)** Aβ(1–28) and **(B)** Aβ(1–28) NoH with and without LMWH. The peptides were incubated without LMWH for 2 hours. After 2 hours, the LMWH was added, and the CD was measured and remeasured 24h after the addition of LMWH.

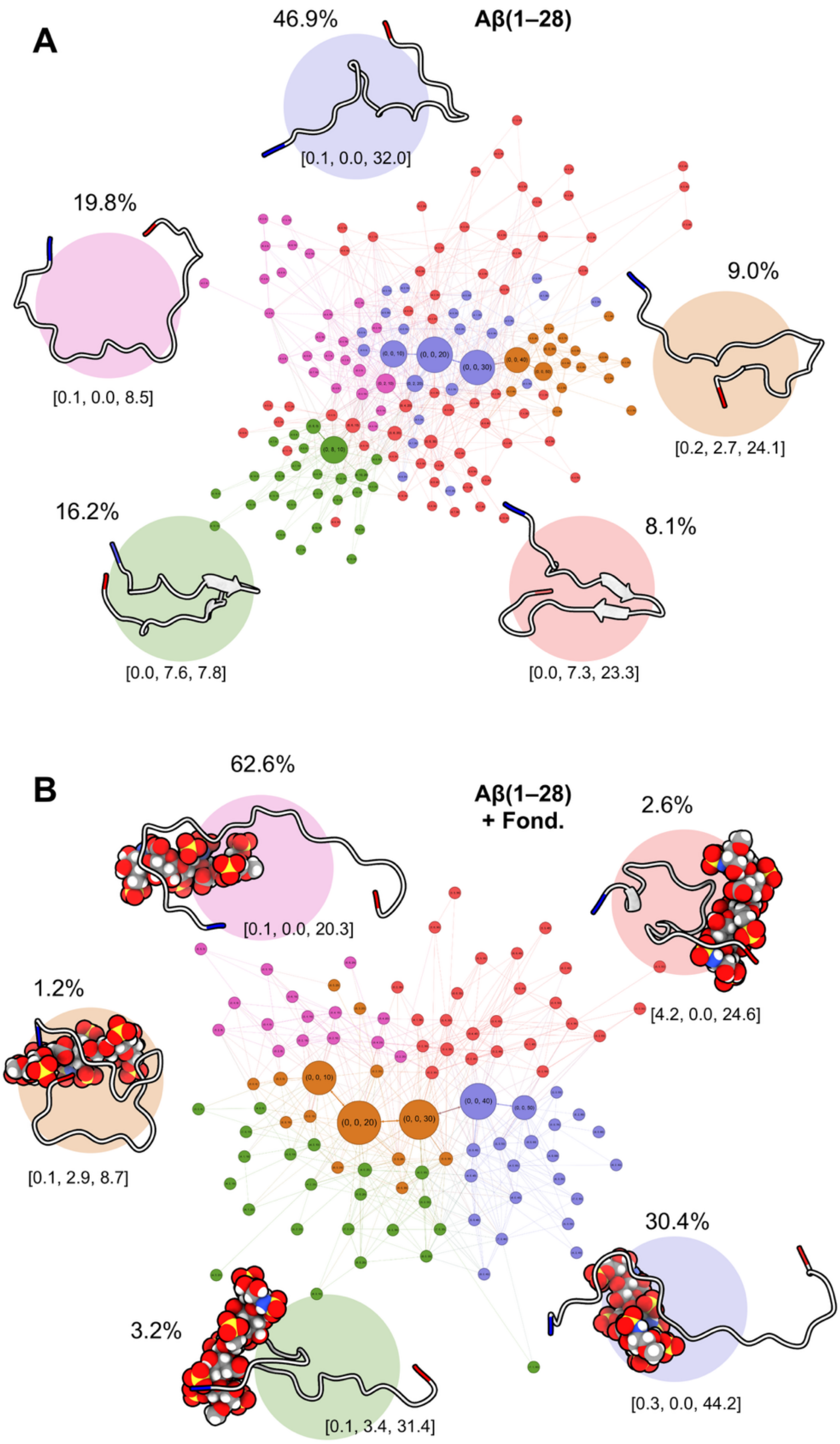

**Figure S10.** Transition network for Aβ(1–28) monomers alone **(A)** and in the presence of equimolar amounts of the pentasaccharide fondaparinux **(B)**. The relative abundance of each conformational cluster is given in percentage. The properties of each cluster are given by [α-helix structure, β-sheet structure, N–C distance].

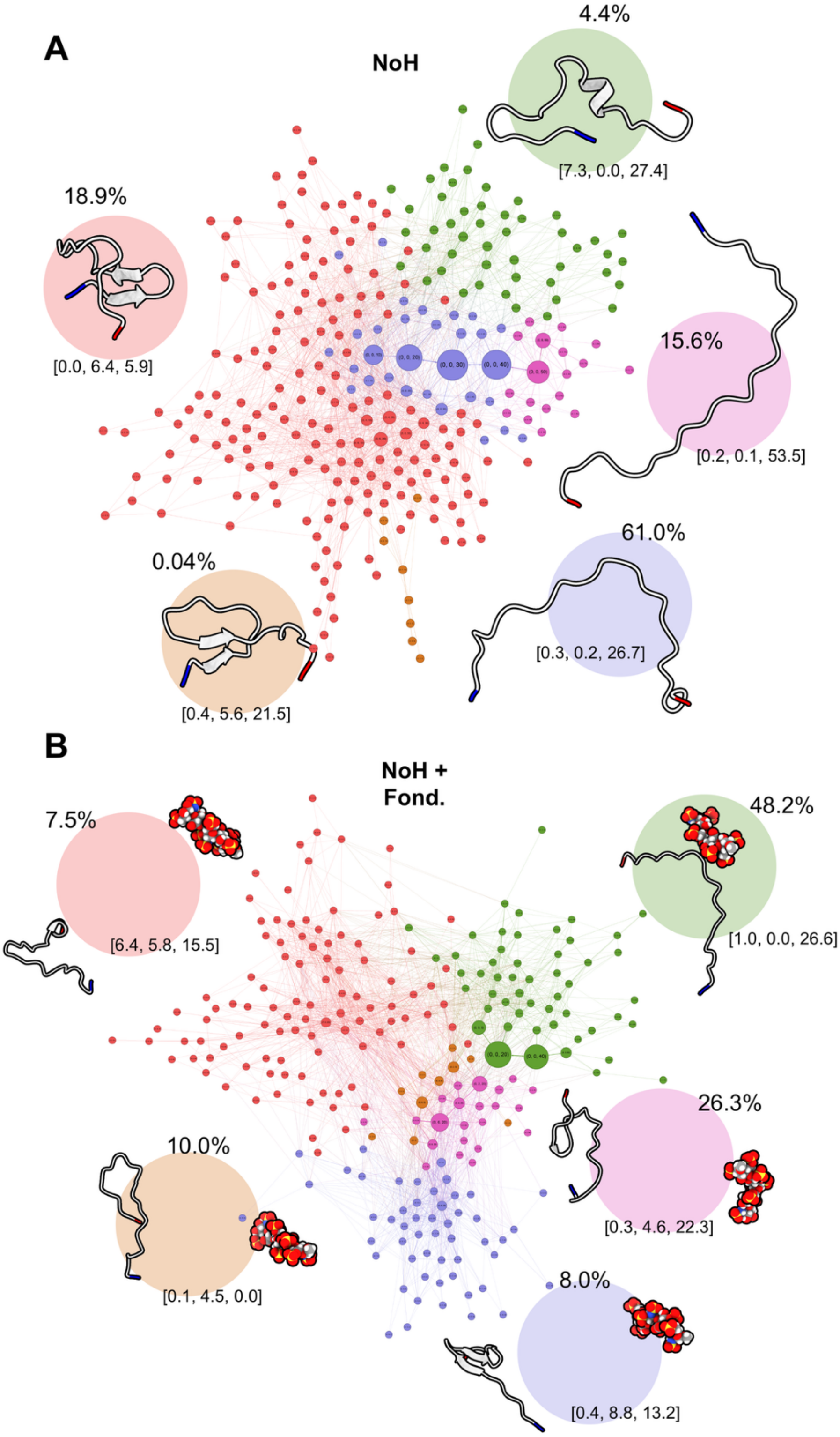

**Figure S11.** Transition network for NoH monomers alone **(A)** and in the presence of equimolar amounts of the pentasaccharide fondaparinux **(B)**. The relative abundance of each conformational cluster is given in percentage. The properties of each cluster are given by [α-helix structure, β-sheet structure, N–C distance].

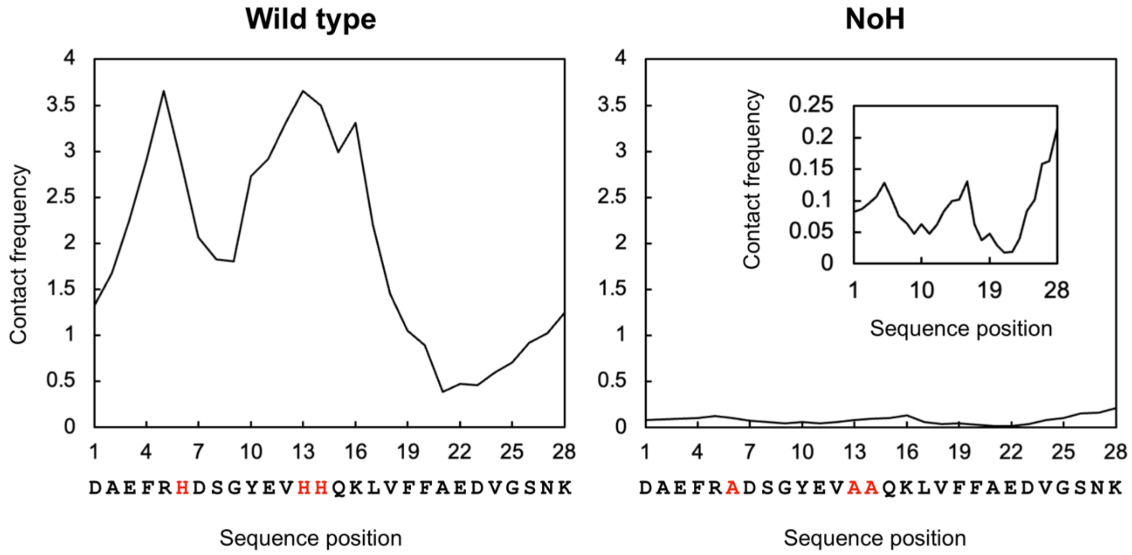

**Figure S12.** Absolute GAG–peptide contact frequencies for Fondaparinux with **(A)** wild-type Aβ(1–28) and **(B)** NoH

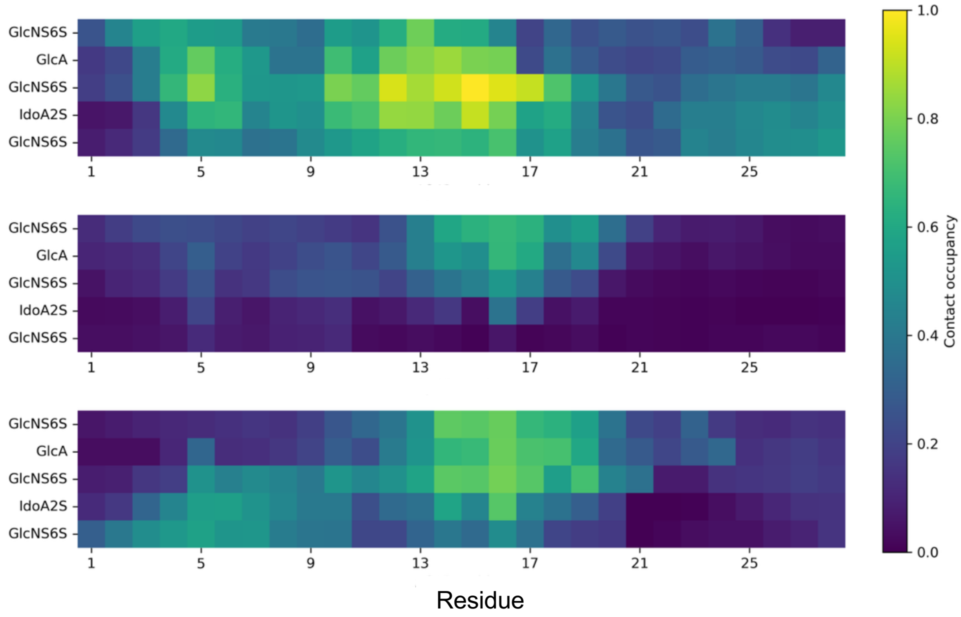

**Figure S13.** Intermolecular GAG–peptide contact maps for Aβ(1–28) trimers in the presence of a single pentasaccharide fondaparinux. The relative contact frequencies for the three peptides are shown separately.

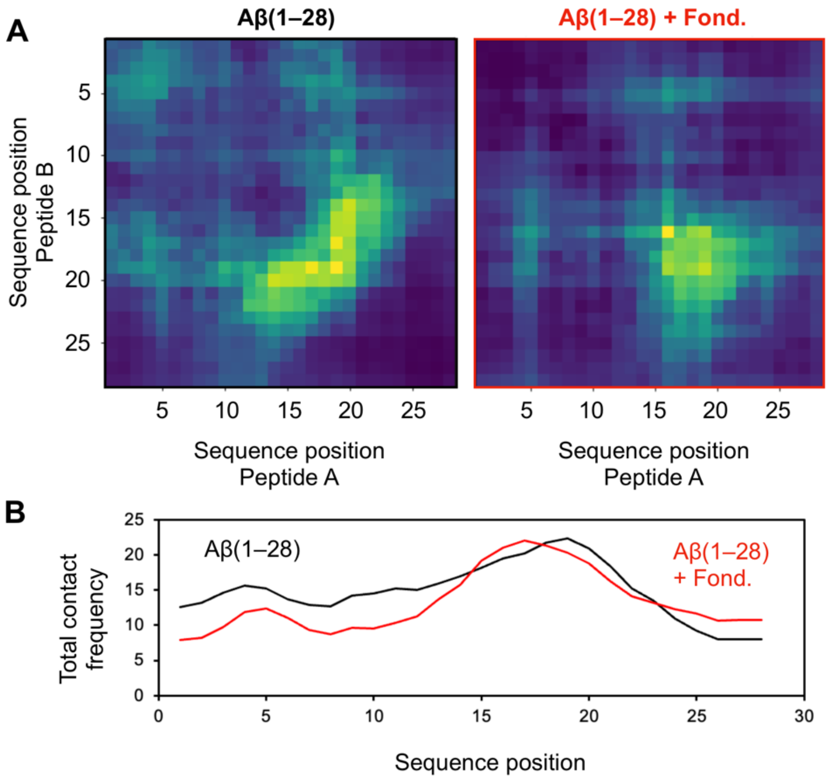

**Figure S14. (A)** Intermolecular peptide–peptide contact maps for Aβ(1–28) trimers alone (left) and in the presence of a single pentasaccharide fondaparinux (right). **(B)** 1D absolute contact frequency profile for Aβ(1–28) trimers alone (black) and in the presence of a single pentasaccharide fondaparinux (red).

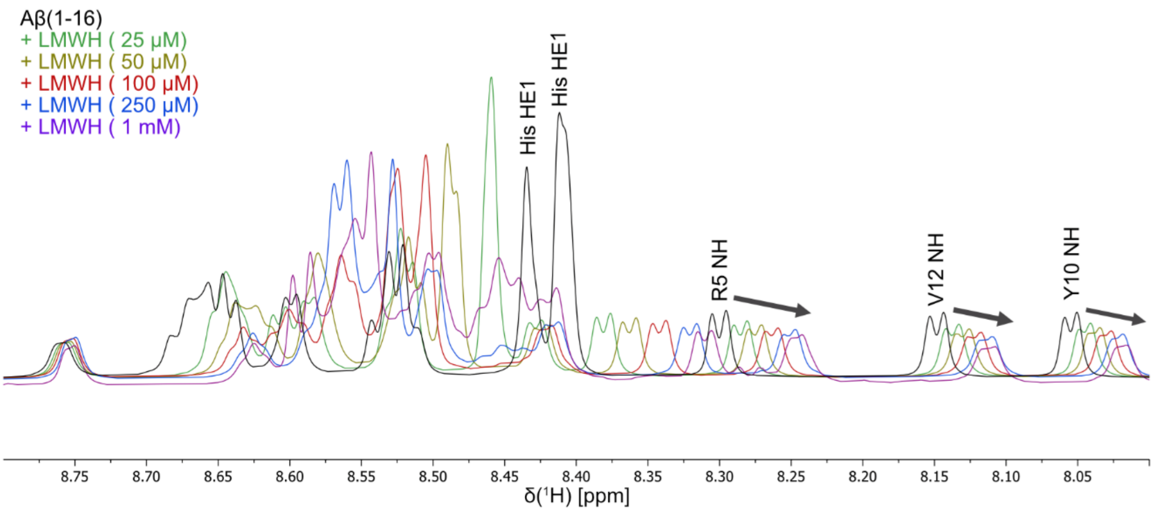

**Figure S15.** $^1$H NMR of 50 μM Aβ(1–16) with increasing concentration of LMWH from 25 μM to 1 mM in 20 mM sodium phosphate buffer pH 6.0.

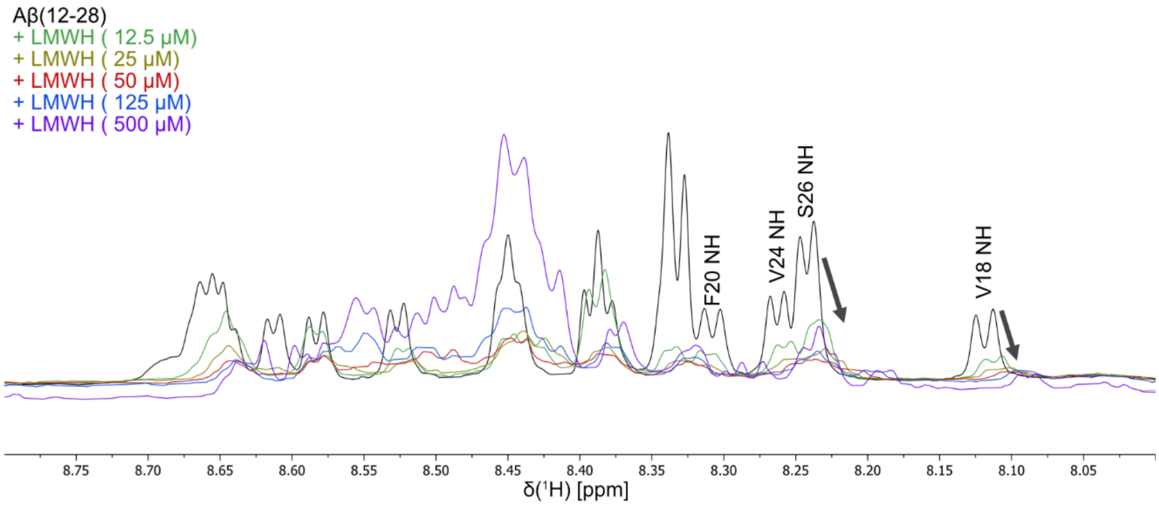

**Figure S16.** $^1$H NMR of 25 μM Aβ(12–28) with increasing concentration of LMWH from 12.5 μM to 500 μM in 20 mM sodium phosphate buffer pH 6.0.

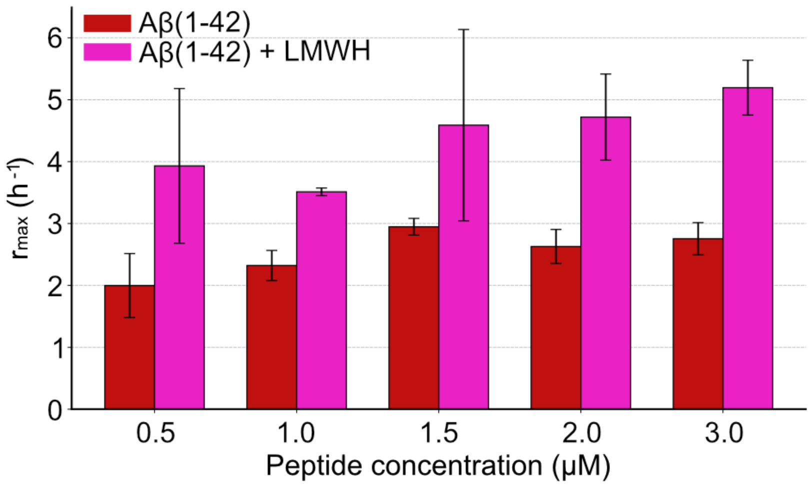

**Figure S17.** $r_{max}$ against concentration plot of data from Figure 4A and B.

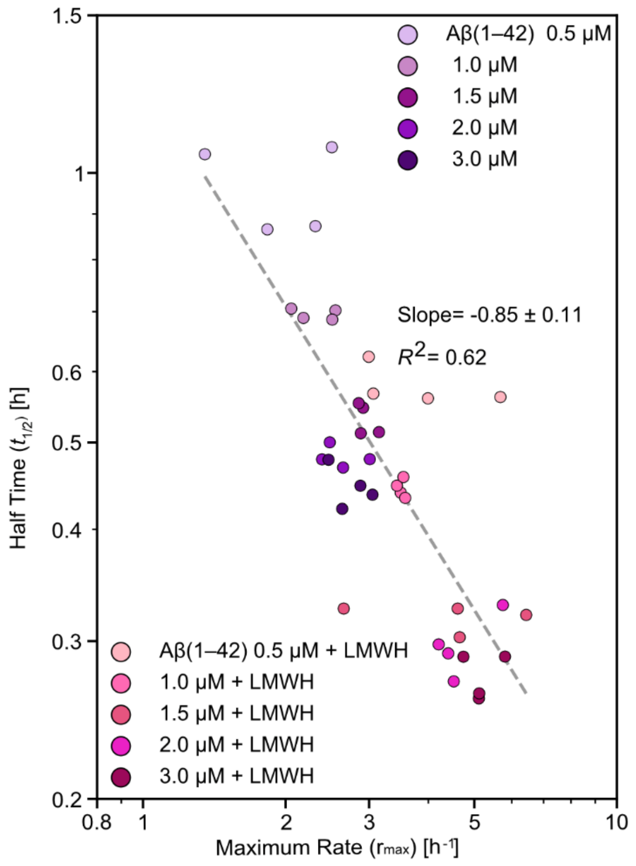

**Figure S18**. $r_{max}$ against $t_{1/2}$ plot from dataset shown in Figure 4A,B. The trend across Aβ(1–42) and Aβ(1–42) + LMWH is similar, resulting in the linear fit slope value close to -1. Aβ(1–42) with LMWH shows overall shorter $t_{1/2}$ and larger $r_{max}$ compared to the Aβ(1–42) only data.

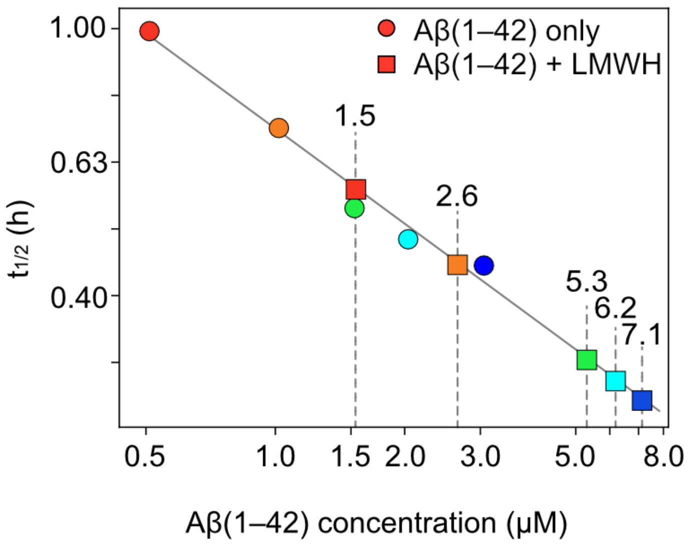

**Figure S19.** Data from Figure 4C fitted to the linear fit of Aβ(1–42) only data. This shows that the half time resulting from the presence of LMWH is similar to nearly threefold increase in the actual concentration of the Aβ(1–42).

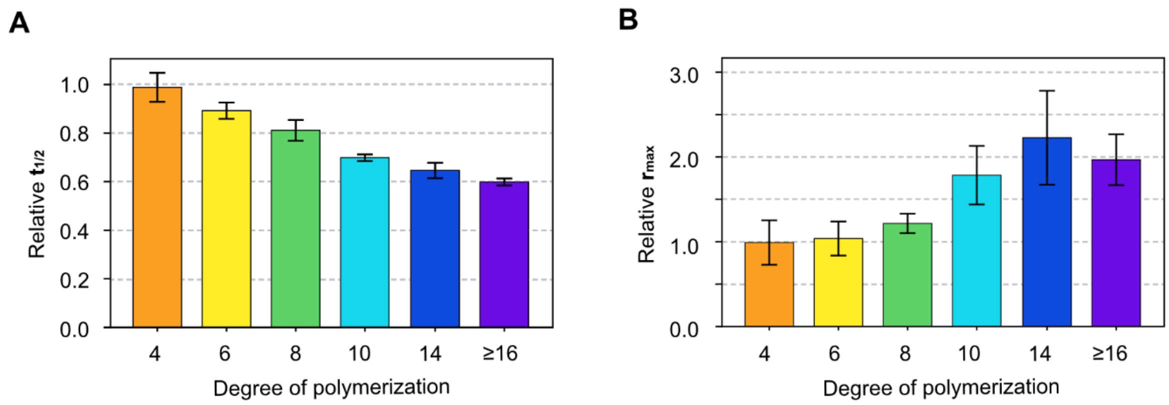

**Figure S20.** Fitted parameter from Figure 5B **(A)** . The relative halftime ($t_{1/2}$) comparison across the degree of polymerization. The results are normalized with the $t_{1/2}$ of Aβ only result as 1.0 **(B)** Relative $r_{max}$ comparison across degrees of polymerization. The results are normalized with the $r_{max}$ of the Aβ-only result as 1.0.

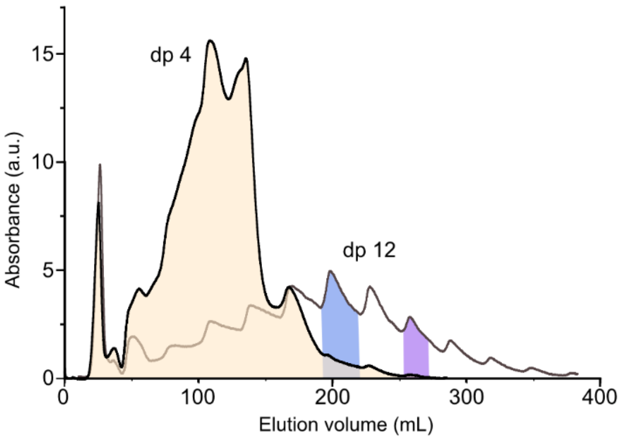

**Figure S21.** Comparison of anion exchange chromatograms of LMWH dp4 and dp12. Overlay of dp4 and dp12 chromatograms demonstrating chain-length-dependent elution. The distinct retention times show that the different chain lengths of LMWH are separated based on their net charge.

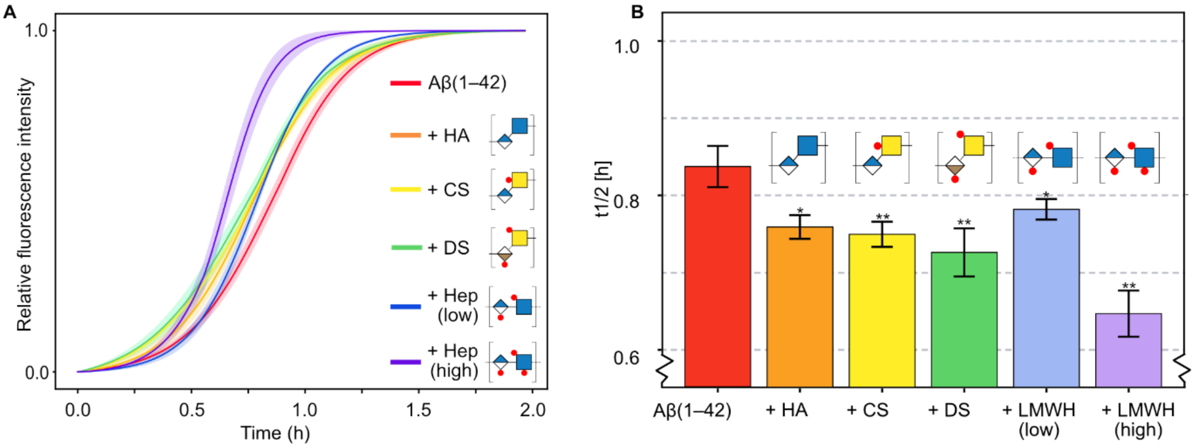

**Figure S22.** The Th T experiment with different GAG classes, hyaluronic acid (HA), chondroitin sulfate A (CS), dermatan sulfate (DS) and heparin (Hep) **(A)** ThT traces for interactions of Aβ(1-42) with different GAG classes (dp12)**.** Equimolar additions of GAGs are incubated with Aβ(1-42) in a 20 mM sodium phosphate buffer pH 6.0. Data are presented as mean ± standard deviation (n = 3). **(B)** The half-time of aggregation from (A) is compared as a bar graph. The significance of each GAG result is compared to the Aβ(1–42) only control.
